# Ascending nociceptive pathways drive rapid escape and sustained avoidance in adult *Drosophila*

**DOI:** 10.1101/2025.10.28.684868

**Authors:** Jessica M. Jones, Anne Sustar, Akira Mamiya, Sarah Walling-Bell, Grant M. Chou, Andrew P. Cook, John C. Tuthill

**Affiliations:** Department of Neurobiology and Biophysics, University of Washington, Seattle, Washington

## Abstract

Nociception — the detection of harmful stimuli by the nervous system — contributes to both rapid escape and long-term avoidance behaviors. *Drosophila* larvae detect damaging heat, mechanical, and chemical stimuli with specialized multidendritic (md) neurons, and these cells are among the only sensory neurons that survive metamorphosis. However, it remains unknown which somatosensory neurons contribute to nociception in adult flies. In an optogenetic screen, we found that abdominal md neurons were the only somatosensory class to induce rapid escape and sustained place avoidance. Calcium imaging from abdominal md axons revealed that they are activated by thermal nociceptive stimuli (>40°C). Connectomic reconstruction showed that md axons form their strongest synaptic connections with ascending interneurons that project to the brain. Among these, we identified two classes of ascending neurons that mediate rapid escape responses and a third that supports sustained avoidance. Our findings reveal that adult *Drosophila* meet several core criteria commonly used to define pain: dedicated nociceptors, ascending pathways connecting peripheral sensors to integrative brain centers, and a capacity for sustained avoidance of noxious stimuli.

## Introduction

The ability to detect and respond to harmful stimuli is a core survival function shared across the animal kingdom. This capacity, known as nociception, relies on specialized sensory neurons called nociceptors, which detect mechanical, thermal, or chemical stimuli with the potential to damage the body^1^. Nociceptor activation triggers fast, reflexive motor programs that help animals avoid injury. In some animals, such as humans, nociceptive responses also contribute to the perception of affective pain, the internal emotional experience that arises from noxious stimuli. Affective pain can motivate long-term behavioral changes, such as learning to avoid particular situations or stimuli^2^.

Whether insects and other invertebrates experience pain has been debated for centuries^3^. Traditionally, insect behaviors were viewed as purely reflexive and lacking an affective component^4^. However, mounting evidence has demonstrated that insects exhibit hallmarks of affective pain, such as learned avoidance^5,6^, behavioral prioritization following noxious stimuli^7^, affective cognition^8^, and sensory generalization across threat modalities^9^. These studies have shown that insects are capable of adapting their behavior to avoid nociceptive stimuli^10^, one of the criteria for affective pain in vertebrate animal models^11–14^. Understanding the neural bases of reflexive and affective responses to nociceptive stimuli in insects could provide fundamental insights into the mechanisms and evolution of pain.

The fruit fly, *Drosophila melanogaster*, offers unique advantages for investigating these questions, including a compact, well-mapped nervous system^15–20^ and powerful genetic tools to enable precise manipulation of specific cell-types. Nociception in *Drosophila* larvae is mediated by class IV dendritic arborization (da) neurons that tile the body wall and trigger escape responses to damaging mechanical, thermal, and ultraviolet stimuli^21^. Class IV da neurons are multidendritic, polymodal nociceptors that express conserved transduction channels including *TrpA1*^22,23^, *painless*^24^, *ppk*^25^, and *Piezo*^26^. Analyses of downstream neural circuits in the fly larva have identified ascending^27,28^ and descending^29^ pathways that relay information from class IV da neurons to higher brain centers and motor neurons, respectively, to coordinate escape behavior^30,31^.

The transformation from larval to adult fly presents new challenges for the somatosensory system^32^. Adult flies possess a different body plan, and their increased mobility exposes them to different threats than ground-dwelling larvae. They also exhibit an expanded behavioral repertoire, including walking, jumping, grooming, courtship, mating, and flight. The somatosensory system of the adult fly is comprised of multiple classes of sensory neurons, including multidendritic neurons, bristles, chordotonal neurons, and campaniform sensilla (**Figure 1A)**, which serve a range of proprioceptive and exteroceptive functions^33^. Multidendritic neurons— class I (proprioceptors in the larva^34^) and class IV da neurons—are among the few sensory neurons that survive metamorphosis from the larval to adult stage, though they undergo extensive remodeling^35–38^. Class I neurons undergo programmed cell death within a week after eclosion, whereas class IV da neurons persist throughout adulthood^39,40^. During metamorphosis, the class IV da neurons are pruned and regenerated into lattice-like dendritic fields^39,40^ that are distributed across the surface of the adult fly’s abdomen, where they are positioned to detect external threats to the abdominal cuticle^41^, similar to free-nerve endings in mammalian skin^42^. (Because the class I–IV nomenclature is not used in the adult fly, we refer to the surviving class IV da neurons as abdominal multidendritic (md) neurons). However, compared to other classes of somatosensory neurons, little is known about the sensory properties of abdominal md neurons, their downstream connectivity, and the behaviors they control.

**Figure 1.**
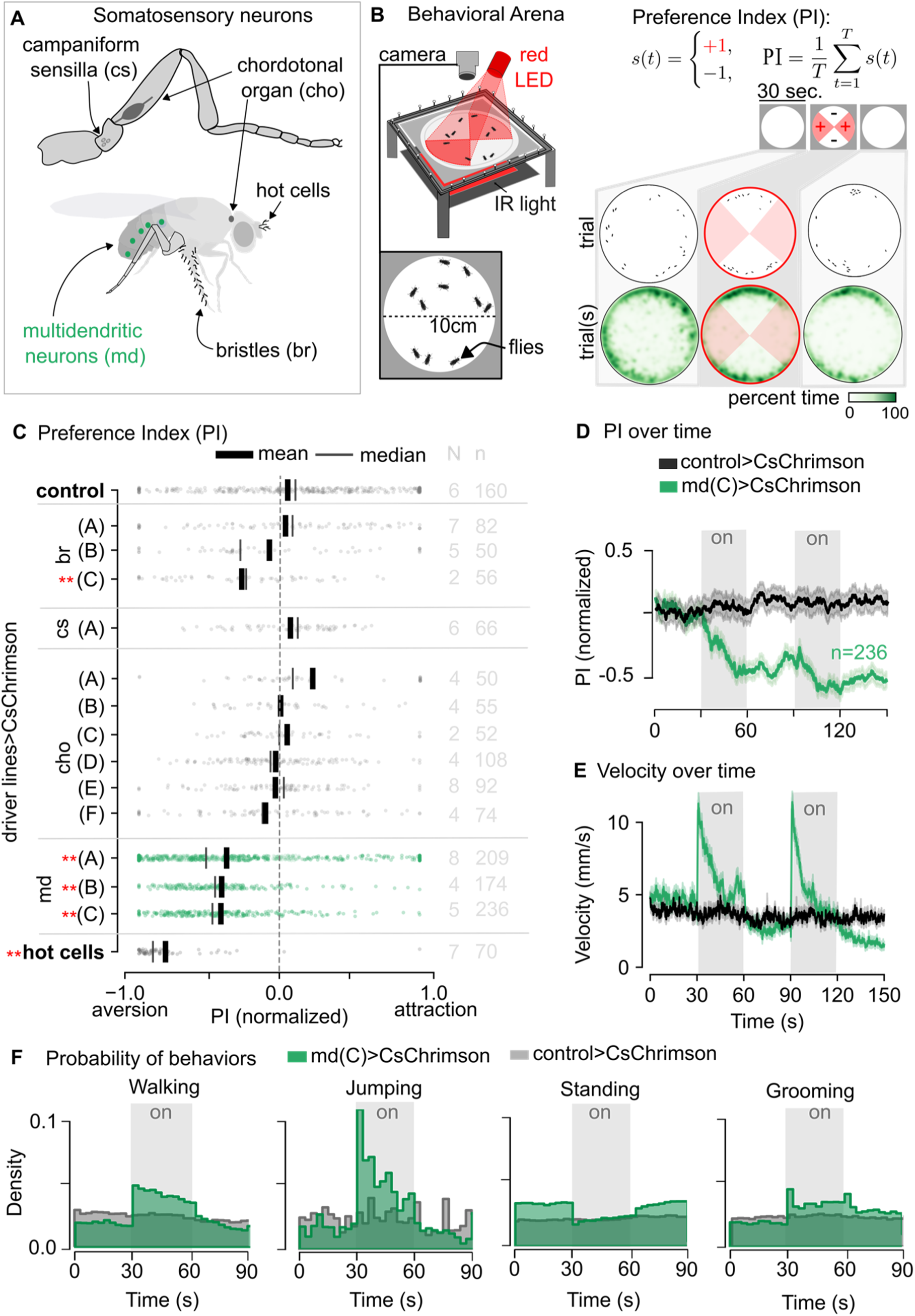
Activation of multidendritic (md) sensory neurons drives place aversion and changes in locomotion. (A) We tested classes of somatosensory neurons in an optogenetic screen. (B) Flies were placed in a 10 cm diameter bowl and their behavior was recorded under infrared light. A red LED delivered optogenetic stimulation during ON trials, and a preference index (PI) was calculated for each fly. Spatial density of flies by number across trials is plotted in green as the percent of time occupying a given region of the arena. (C) Preference indices are shown for different somatosensory neuron classes. The number of trials (N) and flies (n) are indicated for each class. The color scale represents the normalized attraction–aversion index, ranging from –1.0 to +1.0. Letters denote distinct lines labeling subsets of sensory neurons (see Methods for genotypes; md driver line expression is shown in Figure S1). Control flies were of a similar genetic background but lacked Chrimson expression (40B01-GAL4AD >CsChrimson). Red stars mark significant differences relative to controls. Box plots display median, mean, and individual data points. (D–E) Mean preference and velocity traces during 30-second optogenetic stimulation periods illustrate distinct responses between control and md(C)>CsChrimson flies. Gray bars indicate LED ON periods. PI measurements are shown for control flies (n=160) and md(C)>CsChrimson flies (n=236) across multiple trials. PI values were normalized from +1 (attraction) to -1 (aversion). (F) Behavioral probability distributions are plotted over time during 90-second recording periods. Bar heights were normalized within each group to sum to 1.

Here, we investigate the cells and circuits that mediate nociception in adult *Drosophila*. We begin with an optogenetic screen of somatosensory neuron classes, which reveals that abdominal md neurons are the only class that triggers both rapid escape and sustained avoidance. Using *in vivo* calcium imaging, we find that md neurons detect nociceptive thermal stimuli. By combining optogenetics, behavioral analysis, and connectomic reconstruction, we map the central pathways downstream of md axons and identify ascending interneurons that contribute to escape and sustained avoidance. Overall, our results suggest that md neurons in the adult fly abdomen detect nociceptive stimuli and trigger behaviors that are consistent with the experience of pain.

## Results

### A screen for somatosensory neurons that produce avoidance behavior

In some animals, nociceptive stimuli produce both rapid, reflexive escape responses and long-term avoidance. To systematically identify which somatosensory neuron classes produce these behaviors in adult flies, we conducted an optogenetic screen targeting genetically defined populations of somatosensory neurons in adult *Drosophila*^33^. We expressed the red-light activated channelrhodopsin Chrimson in distinct populations of somatosensory neurons and used a four-quadrant assay to quantify behavioral preference over time **(Figure 1A–B).** The fly’s preference or aversion for stimulated regions of the arena served as a readout of stimulus valence, an approach analogous to affective pain assays in mammals^12^.

Most classes of somatosensory neurons, including proprioceptors and touch receptors, elicited only transient behavioral effects, such as grooming or pausing, without producing sustained place avoidance **(Figure 1C)**. However, activation of abdominal md neurons labeled by *ppk*-GAL4 (the md(C) driver line) induced significant and sustained place avoidance **(Figure 1D).** Flies continued to avoid the stimulated region for at least 30 seconds after stimulus offset **(Figure 1D)**, with only a slight decrease in walking velocity (**Figure 1E)**. We confirmed these results in three driver lines that label abdominal md neurons, which produced similar avoidance responses. The magnitude of avoidance increased across consecutive trials of md neuron stimulation (**Figure 1D**). As a positive control, we activated thermosensory hot cells in the antenna, which have been previously shown to produce robust place avoidance in a similar experimental assay^9,43^. Activation of hot cells produced significantly greater place avoidance than md neurons in our experimental setup (**Figure 1C**).

We further classified the behavior of flies in the arena and observed that activating md neurons led to increases in jumping and walking velocity **(Figure 1E–F)**, escape behaviors that also occur in response to other aversive stimuli^44,45^. We noticed that walking velocity peaked then decayed rapidly after stimulus onset, so we further quantified these kinetics using repeated optogenetic stimulation of md neurons in a uniformly illuminated arena **(Figure S1A)**. We found that flies exhibited phase-locked increases in walking velocity even at higher stimulation frequencies (e.g., 2 Hz). Fast walking in response to optogenetic stimulation of md neurons was significantly straighter than normal walking **(Figure S1C–E)**. Overall, our results show that optogenetic activation of md neurons drives both reflexive escape behaviors (jumping and running) and sustained place avoidance.

### Spatially targeted activation of abdominal md neurons triggers stereotyped escape

To resolve the motor responses evoked by md neuron activation with higher temporal and spatial precision, we used optogenetic stimulation of md neurons in tethered flies walking on a spherical treadmill while we tracked the legs and abdomen in 3D **(Figure 2A, Figure S2A–B)**. In addition to enabling high fidelity 3D pose estimation, the tethered preparation allowed us to spatially target md neurons on the abdomen with a laser, thus excluding contributions from other cells labeled by the md GAL4 lines (**Figure S1**).

**Figure 2.**
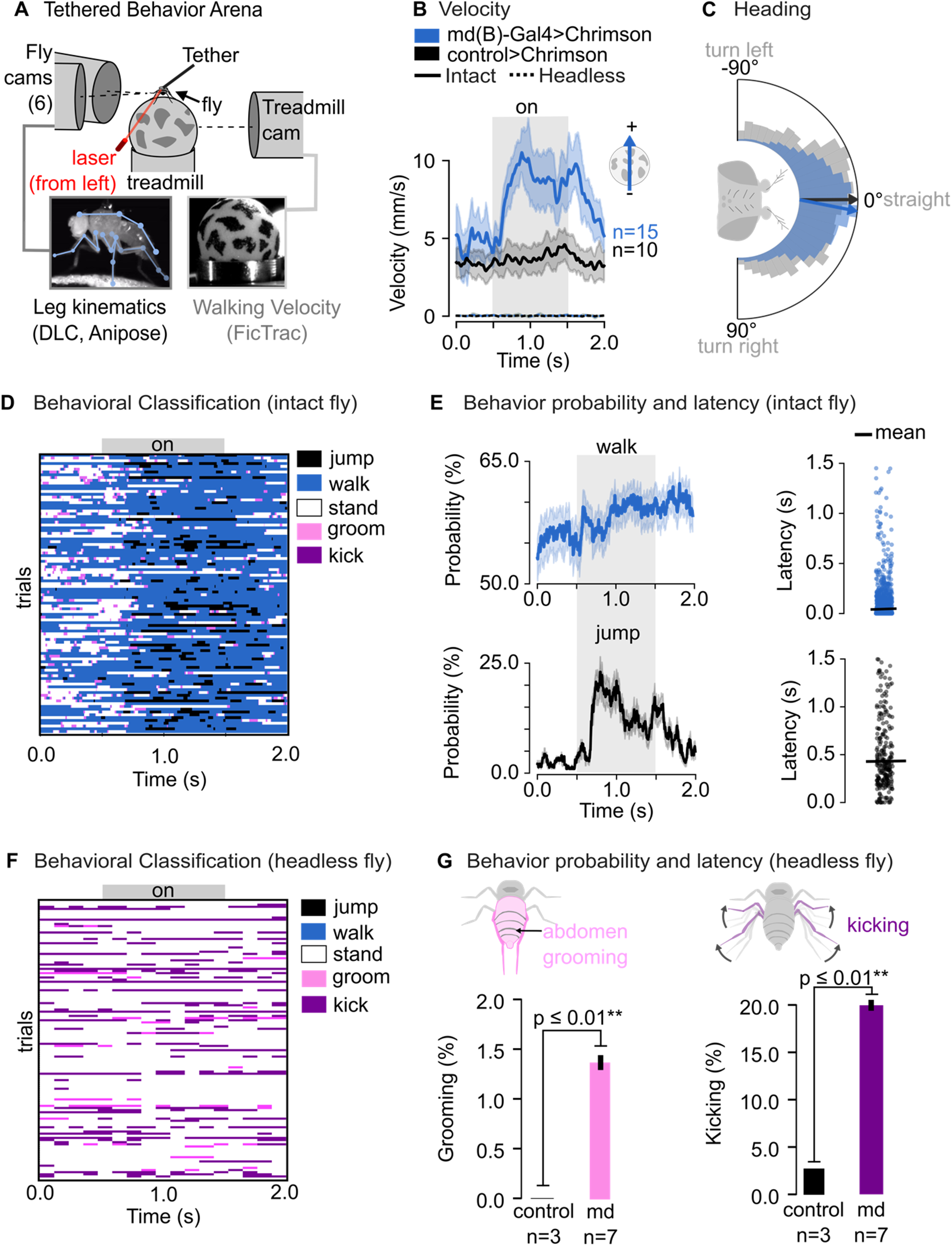
Md neuron activation drives escape behaviors that are absent in headless flies. (A) Flies were mounted on an air-supported spherical treadmill while seven cameras recorded fly behavior and treadmill movement. (B) Average velocity traces (±SEM) are shown for intact flies expressing CsChrimson in md(B) neurons (md(B)>CsChrimson) compared with controls (52A01-GAL4DBD>CsChrimson). Optogenetic activation increased walking velocity. (C) Heading distributions are plotted for control flies (gray) and md neuron–activated flies (blue) during stimulation. (D–E) Behavioral classifications of intact flies revealed transient increases in walking and jumping following stimulation. Panel D shows trial-wise classifications across flies, while panel E plots the mean probability of each behavior over time across flies. (F–G) Behavioral analysis was also performed in headless flies. Ethograms from individual flies across multiple trials (F) illustrate behaviors including walking, jumping, standing, grooming, and kicking. Quantitative probability analyses (G) Trial-wise classifications and mean behavior probabilities between groups compare controls (black) with kicking (purple) and abdomen grooming (pink). Statistical comparisons were made using independent t-tests. All behavioral classifications were derived from automated tracking algorithms. Detailed experimental procedures are provided in the Methods.

Transient (1 sec), spatially targeted (∼350 μm) optogenetic stimulation of md neurons on the left side of the abdomen produced a rapid and robust increase in forward, but not rotational, velocity. Forward velocity increased to twice the baseline level within 500 ms of stimulus onset **(Figure 2B),** though the flies’ heading angle did not significantly change **(Figure 2C).** This velocity increase was frequently accompanied by jumping behavior, during which the fly released the ball and retracted its leg **(Figure 2D–E).** Increases in walking speed occurred at a shorter latency (mean=100 ms) than jumping (mean=480 ms; **Figure 2E).**

Previous work has shown that flies and other insects exhibit some sensorimotor reflexes in the absence of descending input from the brain (i.e., in decapitated flies^46^). Decapitated flies may also jump^47^ and attempt coordinated walking^48^ during thermo- or optogenetic stimulation of specific neurons. To determine whether md neuron-mediated escape responses require the brain, we repeated our experiments in decapitated flies. While intact flies exhibited coordinated escape behaviors, such as increased forward locomotion and jumping, we found that decapitated flies performed leg kicking and site-specific grooming directed at the abdomen. Kicking and grooming were variable but occurred at a significantly higher percentage over the entire stimulus period than controls **(Figure 2F–G)**. Decapitated flies never displayed coordinated walking on the treadmill **(Figure 2F)**, indicating that ascending or descending projections to/from the brain are essential for escape initiation following md neuron activation. Interestingly, a study in mice also found that brain circuits are required to coordinate escape behavior^49^.

Our findings collectively indicate that activation of abdominal md neurons in adult *Drosophila* drives escape responses: jumping and forward locomotion. The same stimuli in headless flies produced distinct motor reflexes: kicking and abdominal grooming. This suggests that nociceptor-mediated escape behaviors rely on ascending and descending signals to and from the central brain.

### Abdominal md neurons are sensitive to noxious heat

Knowing that optogenetic activation of md neurons produces escape and sustained avoidance, we sought to understand the stimuli that they sense. The axons of md neurons project into the abdominal ganglion, the most posterior compartment of the fly ventral nerve cord (VNC). To characterize the sensory response properties of md neurons, we recorded their axonal calcium activity in the abdominal ganglion with *in vivo* two-photon imaging **(Figure 3A).** We expressed GCaMP7f and tdTomato in md neurons and quantified their activity as the ratio of green to red fluorescence while applying thermal and mechanical stimuli that had previously been shown to evoke class IV da neuron activity in larval *Drosophila*^50–53^. Gentle deflection of the abdomen with a 40°C metal probe elicited robust calcium responses in md axons (**Figure 3B)**. Elevated calcium levels were sustained throughout the duration of the stimulus **(Figure 3B–D).** We also observed increased calcium activity when we topically administered AITC, an agonist of Trp channels including *TrpA1*^54^, to the abdominal surface **(Figure S3)**. This result suggests that *TrpA1* contributes to the detection of noxious thermal stimuli, as has been shown in *Drosophila* larvae^55^. Notably, we did not observe consistent mechanosensory responses when a room temperature (25°C) probe touched the abdomen (**Figure 3B)**. Overall, our results show that abdominal md neurons in adult *Drosophila* detect noxious heat **(Figure 3B–D)**, responding at the same high temperature (40°C) previously used in fly larvae^22,56^.

**Figure 3.**
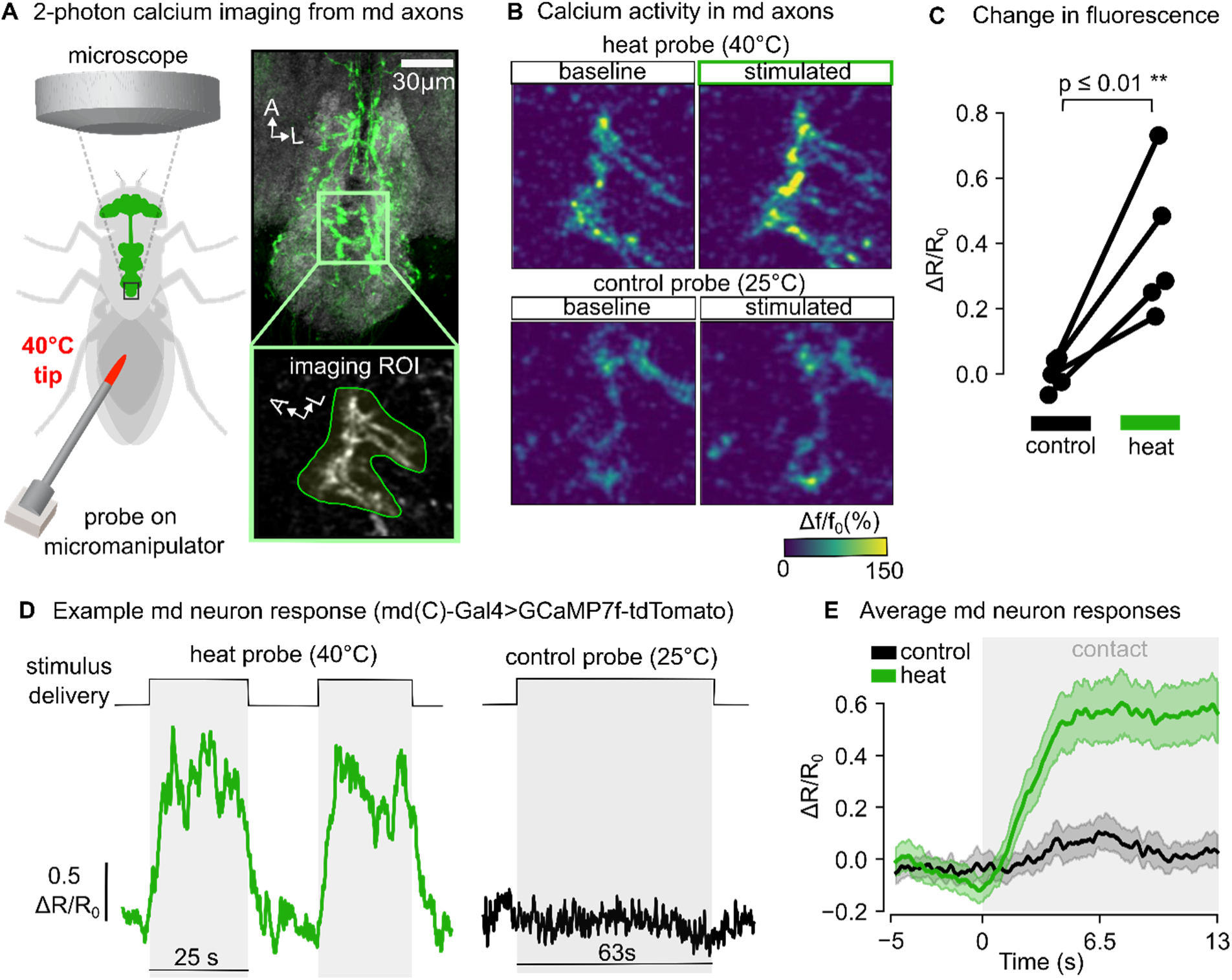
Calcium imaging from md axons reveals sensitivity to high temperature. (A) Two-photon calcium imaging from md axons in the abdominal ganglion. The schematic illustrates the imaging configuration with a temperature probe for thermal stimulation (left). A confocal image displays md neurons expressing GFP (green; md(C)-GAL4>UAS-GFP) within the ventral nerve cord, with the region of interest (ROI) outlined (scale bar = 30 μm). The ROI image shows GCaMP7f expression at md axon terminals (md(C)-GAL4>UAS-GCaMP7f-tdTomato). (B) Calcium responses of md neurons are compared under different temperature conditions: a 25°C probe (room temperature control) and a 40°C probe (heat stimulation). (C) Response magnitudes are quantified across flies, with ratio values plotted for each fly (n=5). All flies exhibited substantial increases in calcium responses during heat stimulation, with values ranging from ∼0.2 to 0.8. (D) Representative GCaMP7f traces from md axons in a single fly are shown. The left trace captures calcium transients during 40°C probe stimulation over a 25-second recording period, while the right trace shows minimal activity (ΔR/R_0_ = 0.5 scale) during room-temperature probe application over 63 seconds. Stimulus timing is indicated by gray bars above each trace. (E) Average calcium responses over time are plotted, with the green trace representing mean ± SEM responses to the 40°C probe and black to the 25°C probe. Calcium levels remained elevated (∼0.6 ΔR/R_0_) throughout the stimulation period (gray shading).

### The axons of nociceptive md neurons form a somatotopic map in the abdominal ganglion

We next analyzed the spatial organization of md neuron axons in the abdominal ganglion using genetic labeling, light microscopy, and connectomics **(Figure 4A)**. Light microscopy images of GAL4 lines labeling md axons revealed 30 md neurons with extensive dendritic arbors distributed across the fly abdomen (**Figure S4**). We counted 16 ventral and 14 dorsal md neurons per fly, each spanning one tergite or sternite on either side. Md axons enter the abdominal ganglion via four different nerves and then project ventrally within the neuropil. For example, nerve 2 carries four axons (two per side) from segment 1, nerve 3 carries four axons, and the nerve trunk carries six from segments 6/7, with nerve 4 carrying the remaining 15.

**Figure 4.**
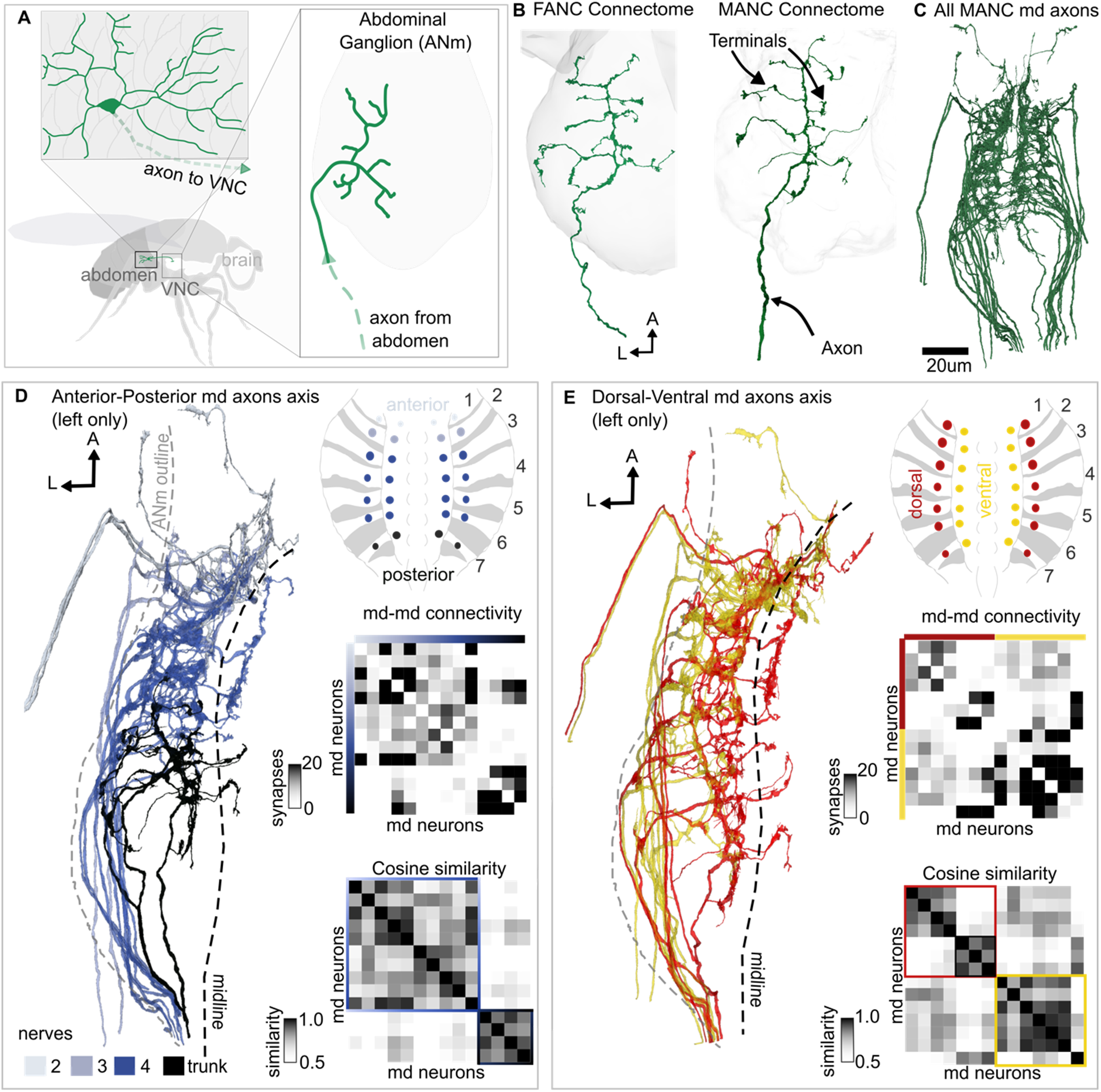
Axons of abdominal md neurons form somatotopically organized and stereotyped terminal arbors in the abdominal ganglion. (A) A schematic of the abdominal ganglion shows axonal projections from peripheral md neurons. Individual axons (green) arborize in stereotyped patterns within the neuropil (grey outline). (B) An anatomical overview depicts md neuron organization within the Female Adult Nerve Cord (FANC) and Male Adult Nerve Cord (MANC) connectomes, revealing consistent innervation patterns across datasets. (C) All md neurons reconstructed in MANC are shown (n=30). (D–E) Three-dimensional reconstructions of axon terminals demonstrate spatial organization along the dorsal–ventral and anterior–posterior axes. Axons display biased output distributions depending on peripheral soma position and exhibit like-to-like connectivity patterns when terminals are in close proximity. Pairwise cosine similarity analyses compare axonal arbor morphology (left) and synaptic connectivity (right) for md neurons grouped by dorsal (n=8) versus ventral (n=7) soma location in panel E, and by anterior–posterior position in panel D. Schematics illustrate the spatial organization of md neurons within abdominal segments 1–7. Color coding indicates cell body origins and nerve pathways, with trunk nerves 3 and 4 highlighted. Neurons with similar spatial locations on the abdomen exhibit more similar connectivity patterns.

We used morphological criteria to identify md axons in existing electron microscopy volumes of the female (FANC)^15,18^ and male (MANC)^19^ adult nerve cords (**Figure 4B**). In FANC, we manually proofread each md axon. The axons in MANC were already proofread after automated segmentation. These connectome reconstructions revealed that individual md axons arborize in multiple segments of the abdominal ganglion, with unique projections into each segment. Md axon projections are spatially segregated from abdominal mechanosensory bristle and gustatory axons, as previously described^57^ **(Figure S4A–F).** Md axons in FANC and MANC have qualitatively similar terminal structure, synapse distribution, branching architecture, and input-output connectivity, suggesting that md neurons are similar in male and female flies, despite known sexual dimorphisms in other aspects of fly behavior and neural circuit organization^58^.

To determine the relationship between each md neuron’s peripheral location on the abdomen and their central projection into the abdominal ganglion, we used sparse genetic labeling with SPARC (Sparse Predictive Activity through Recombinase Competition)^59^. Labeling single neurons revealed a somatotopic organization of md axons within the abdominal ganglion (**Figure S4G– M**), which we then used to infer the peripheral origin of each md axon in the connectomes (**Figure 4D–E**). We found that md neurons from anterior abdominal segments terminate in the most anterior region of the abdominal ganglion, while md neurons on the posterior abdomen arborize posteriorly (**Figure 4D**). We also observed somatotopy along the dorsal-ventral axis. The axons of md neurons from the dorsal abdomen cross the midline of the abdominal ganglion before terminating, while ventrally originating neurons remain ipsilateral to the midline **(Figure 4E).** This crossing pattern is similar to the pattern of somatotopy described in the larval nervous system^60^.

We also observed somatotopic structure in the downstream synaptic connectivity of md neurons. Dorsal md neurons are more likely to synapse on the axons of other dorsal neurons, and vice versa for ventral neurons. Dorsal md neurons exhibit more similar postsynaptic connectivity to other dorsal or ventral neurons, as measured by their cosine similarity **(Figure 4E, inset)**. We also observed clusters of downstream connectivity among anterior and posterior axons **(Figure 4D, inset)**. These results reveal that md axons are organized somatotopically in the fly abdominal ganglion, which may facilitate integration of correlated sensory signals by downstream circuits.

### Nociceptive abdominal md axons are most strongly connected to ascending neurons

We next used the connectome to analyze the synaptic inputs and outputs of md neurons. We first analyzed presynaptic input to md axons, i.e., feedback from VNC neurons. All neurons in the MANC connectome have a predicted neurotransmitter, based on a validated machine learning classifier^61^. Unlike other classes of sensory neurons that receive presynaptic inhibition (e.g., leg proprioceptors^62^), we found that md neurons primarily receive cholinergic input, which is typically excitatory in the fly CNS **(Figure S5A–E).** Excitatory feedback to md axons could contribute to sensitization, a hallmark of nociceptors in vertebrates^63,64^ and larval *Drosophila*^65,66^.

We next analyzed the postsynaptic VNC targets of all 30 md axons, focusing on neurons receiving ≥4 synapses from abdominal md neurons, a threshold previously used to filter for functionally relevant connections in MANC^67–69^. We identified 374 distinct postsynaptic neurons that we categorized into six morphological classes: ascending neurons, descending neurons, local interneurons, motor/efferent neurons, and other sensory neurons **(Figure 5A).** We found that md neurons form particularly strong connections onto ascending neurons **(Figure 5B–5D).** Nearly half (49%) of md axon synapses are onto ascending neurons, whereas other somatosensory classes, tactile bristles and leg proprioceptors, have less than 30% of their synapses onto ascending neurons, with the majority onto local neurons **(Figure 5C).** Approximately 73% of the ascending neurons postsynaptic to md axons are cholinergic **(Figure 5E**), compared to 76% for bristles and 54% for proprioceptors. Overall, this connectivity suggests that sensory information from md axons is rapidly routed to the brain, compared to the tactile and proprioceptive systems, which predominantly feed into local VNC circuits.

**Figure 5.**
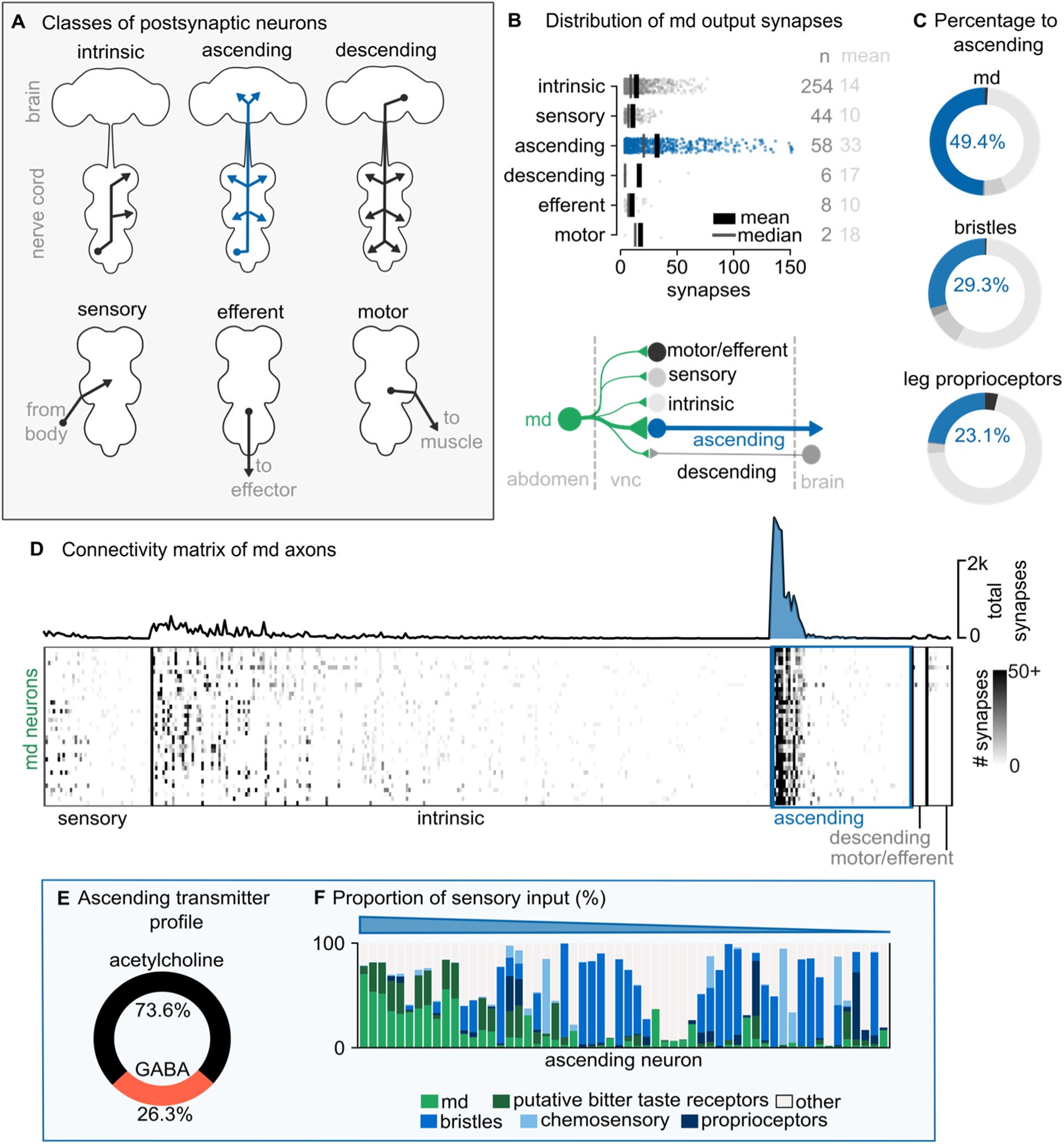
Ascending neurons are the main postsynaptic target of abdominal md sensory axons. (A) Classes of VNC neurons receiving input from md axons. (B) Md connectivity patterns are illustrated at the bottom, highlighting synaptic relationships with intrinsic, sensory, efferent, motor, descending, and ascending neuron types. The top panel quantifies synaptic connectivity by target class, with a strip plot displaying mean synapse counts and individual data points onto postsynaptic VNC neurons. (C) The percentage of synapses to ascending neurons is summarized in a pie chart, with comparisons to mechanosensory bristles and leg proprioceptors. (D) A connectivity matrix reveals strong connections to a small number of ascending neurons. Synapse counts between md neurons and their targets are represented as a heat map, with color intensity corresponding to connection strength (0 to >150 synapses). (E) The predicted neurotransmitter identities of ascending neurons are shown as a pie chart, grouped by transmitter percentage. (F) Sensory input proportions for each postsynaptic ascending neuron.

A small number of ascending interneurons (∼20) receive a disproportionately high number of synapses from md axons. These ascending neurons also receive most of their sensory input from md neurons vs. other sensory neuron classes **(Figure 5F)**, suggesting that these ascending neurons may be specialized for conveying nociceptive signals from abdominal md neurons to the brain.

### Different components of escape and sustained avoidance are mediated by distinct ascending pathways

We sought to identify genetic driver lines that label the primary ascending neurons downstream of md axons. We first searched in the MANC connectome for ascending neurons that received >20% of their input from md neurons. This resulted in 22 individual ascending neurons. We used NeuronBridge^70^ to identify three split-GAL4 driver lines that labeled different anatomical subtypes of ascending neurons that fulfilled these criteria **(Figure 6A).** The ascending neurons labeled by these driver lines all arborize throughout the abdominal, hindleg, and wing neuropils. Within the brain, all the ascending axons arborize within the gnathal ganglion (GNG), a premotor center. However, individual ascending neurons within each driver line project to distinct brain areas, suggesting distinct functions.

**Figure 6.**
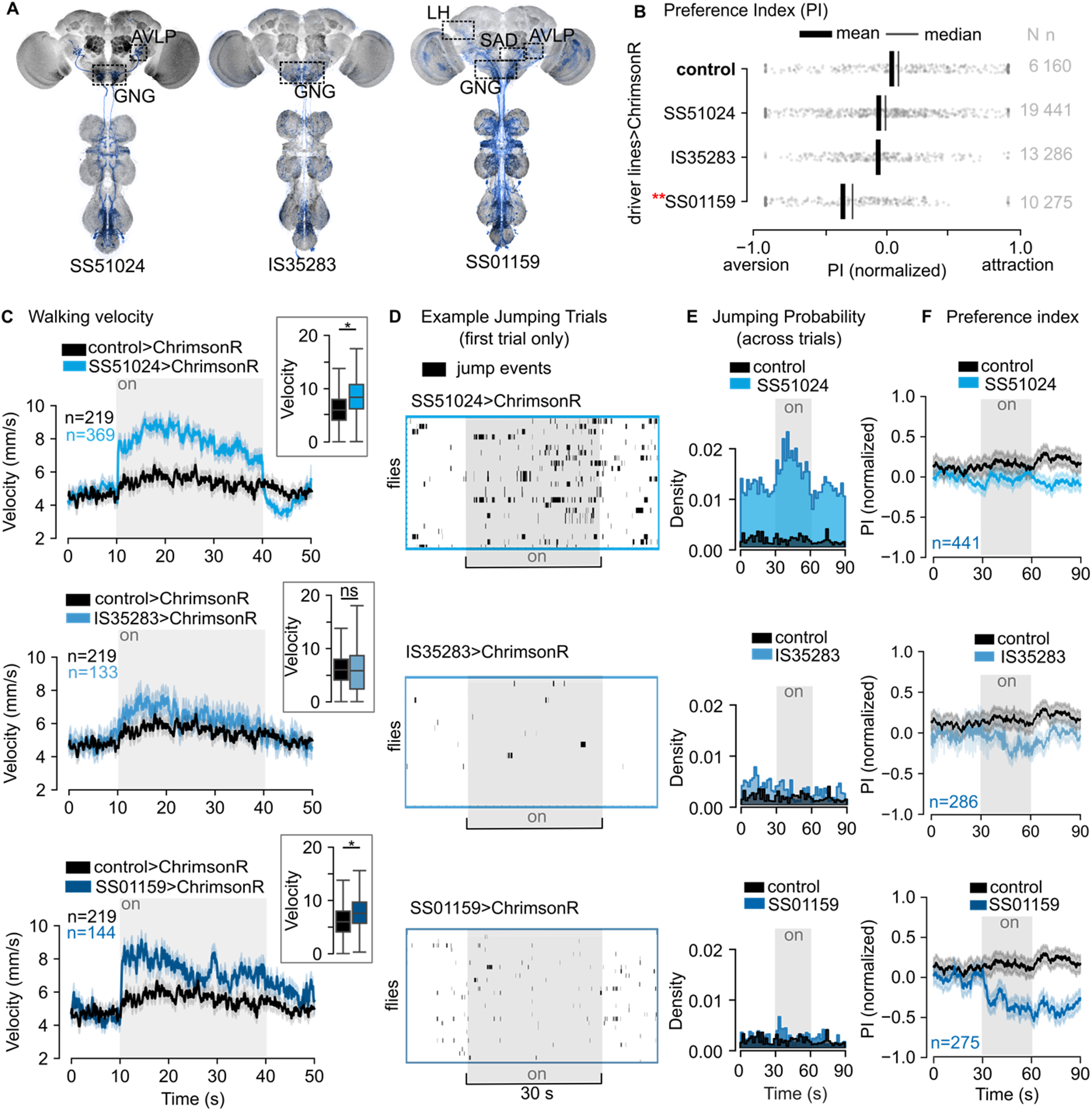
Optogenetic activation of ascending neurons produces escape and sustained avoidance. (A) Schematic shows the brain regions targeted by three genetic lines: SS51024, which projects to the AVLP and GNG regions; IS35283, which projects to the GNG; and SS01159, which projects to the LH, SAD, AVLP, and GNG regions. GFP expression is pseudocolored in blue. (B) The preference index is shown for all second-order driver lines, with a red star denoting statistical significance. Control flies carried the genotype 40B01-GAL4AD>CsChrimson. (C) Mean walking velocity traces are shown over time for control flies (black) and optogenetically stimulated flies (blue) over 50-seconds. The light stimulation period is indicated by grey bars. Sample sizes of flies were SS51024: n=369, IS35283: n=133, and SS01159: n=144 flies. Shaded areas represent the standard error of the mean. Insets show the change in velocity for the first 10 seconds during optogenetic activation between controls in black, and experimental genetic lines in blues. (D) Representative raster plots show jumping events (black bars) during the first trial for individual flies across the stimulation period. Each row represents one fly, with jumping events plotted against time. The light stimulation period (30 seconds) is indicated by grey shading. (E) Population-level histograms show jumping probabilities comparing control (black) and stimulated (blue) conditions across 90-second trial periods. Light stimulation occurred during the middle 30 seconds, indicated by grey shading. (F) Preference indices quantify behavioral responses over time, calculated as the difference between stimulated and control conditions. Statistical comparison to controls during stimulation period quantified in B. Sample sizes were SS51024: n=441 flies, IS35283: n=286, and SS01159: n=275. Values above zero indicate increased preference relative to controls, while values below zero indicate decreased preference. All traces show mean ± SEM. AVLP, anterior ventrolateral protocerebrum; GNG, gnathal ganglion; LH, lateral horn; SAD, saddle

We used optogenetic activation of each split-GAL4 line to test whether they recapitulate the behaviors we observed when activating the md sensory neurons in freely walking flies. Activating all three driver lines produced increases in walking velocity **(Figure 6C).** The line with neurons projecting to the anterior ventrolateral protocerebrum (AVLP; SS51024) was the only one among the three that displayed elevated jumping behavior **(Figure 6D–E).** We also noticed that this line increased jumping outside of the optogenetic stimulus period. The third driver line (SS01159) contained ascending axons that project to the AVLP, saddle (SAD), and lateral horn (LH), a higher-order olfactory region **(Figure 6A).** This driver line labeled a broader set of neurons than the other two but was the only one among the three that produced sustained place avoidance **(Figure 6B, F).** These results are consistent with a recent study showing that activation of this same driver line produced aversive learning in a similar spatial avoidance paradigm^45^. Overall, our results suggest that ascending pathways downstream of md axons drive distinct, though partially overlapping, aspects of rapid escape and sustained avoidance behavior.

Finally, we analyzed the downstream connectivity of the different classes of ascending neurons using existing brain and nerve cord connectomes (MANC^19^, FAFB^67,68^, and BANC^16^). Matching cells labeled by each driver line to ascending neurons in the connectome revealed that they connect to distinct downstream circuits **(Figure S6).** SS51024 neurons are strongly connected to the giant fibers and other premotor networks that produce rapid takeoff responses^71,72^. IS35283 neurons primarily target walking control circuits, including connections to descending neurons that modulate locomotion speed and direction^48,73–75^. SS01159 neurons connect to brain regions associated with learned avoidance, including pathways to dopaminergic clusters that mediate aversive memory formation^76,77^. These anatomical analyses suggest potential circuits through which ascending neurons could transform nociceptive signals from abdominal md neurons into distinct escape and avoidance behaviors.

## Discussion

### Comparison of nociception to other somatosensory modalities

Several features distinguish md neurons from other somatosensory neurons in the adult fly. First, unlike proprioceptors and touch receptors, we found that md neurons drove both rapid escape and sustained place avoidance when optogenetically stimulated **(Figures 1, 2)**. Activating other somatosensory neurons in our optogenetic screen produced only transient effects, such as grooming or pausing, without sustained place avoidance. An exception was the bristle driver line b(C), which produced a significantly negative preference index, though not as low as any of the md driver lines. The b(C) line labels mechanosensory bristles on the eye, and closer inspection of the video revealed that optogenetic stimulation of this line produced sustained head grooming, similar to behavior seen in a previous study using this driver line^78^.

Second, abdominal md neurons exhibit distinct connectivity patterns compared to other somatosensory neurons. While mechanosensory bristles and proprioceptors primarily connect to local motor circuits in the VNC, we found that md neurons dedicate a large fraction of their synaptic output to ascending neurons that project to the brain **(Figure 5)**. This connectivity pattern suggests that nociceptive information requires integration with higher-order circuits for threat assessment and long-term behavioral modification.

We also found that abdominal md neurons have distinct properties from other thermosensory neurons in adult *Drosophila.* Antennal “hot cells” detect warm ambient temperatures (30–35°C) and drive thermotaxis behaviors, such as navigation toward the fly’s preferred temperature^9,22^. In contrast, our results suggest that abdominal md neurons function as nociceptors that detect noxious heat (>35°C) **(Figure 3).** Hot cells detect heat using the gustatory receptor GR28B(D)^79^, while past work in the larva suggests that class IV da neurons rely on TRPA1^23^. The difference in temperature sensitivity between md neurons and hot cells is also reflected in their circuit organization: the ascending neurons downstream of md axons do not converge with lateral horn circuits downstream of antennal hot cells^80,81^. Additionally, hot cells guide turning behaviors that enable flies to navigate thermal gradients^9,22^, whereas we did not find directional turning in response to unilateral md neuron activation. These differences suggest that the fly nervous system uses distinct thermosensory neurons and downstream circuits to detect and respond to innocuous and noxious temperatures.

The somatotopic organization we observed in abdominal md axon terminals **(Figure 4D–E)**—both anterior-posterior and dorsal-ventral organization—is similar to that described in the 3^rd^ instar fly larvae^60^. Our cosine similarity analysis revealed that md neurons from similar abdominal regions share similar downstream connectivity, which could support spatially coordinated responses, though we did not observe directionality in escape responses following unilateral optogenetic stimulation. Analysis of postsynaptic connectivity revealed organizational principles beyond simple somatotopy (**Figure S5F**). Most postsynaptic partners receive input from all md neurons along the anterior-posterior axis, with few neurons receiving selective input from md neurons within the same nerve. However, dorsal and ventral-originating md neurons connect to different postsynaptic cells, with target neurons distributed along the dorsal-ventral axis. Neurons postsynaptic to dorsal md neurons arborize more medially in the VNC than those receiving ventral md input. This asymmetric organization suggests that VNC circuits may preferentially integrate threat information along the body’s anterior-posterior axis while maintaining separate processing channels for dorsal vs. ventral stimuli. This organization could reflect distinct escape strategies in response to nociceptive stimuli from above versus below the fly.

### Comparison of nociceptors across metamorphosis

Insects that undergo complete metamorphosis develop two distinct body forms during their lifetime: a larva specialized for feeding and growth, and an adult built for reproduction and dispersal. This transformation involves dramatic changes to the nervous system, which is extensively reorganized through the differentiation of adult-specific neurons, the programmed death of certain larval neurons, and the structural remodeling of others^82,83^. Multidendritic sensory neurons, including the abdominal md neurons, are among a small minority of identified neurons that are known to survive through metamorphosis. Other da neuron types also survive metamorphosis but are not labeled by the *ppk*-GAL4 lines we used in this study^37^. Multidendritic neurons have been extensively studied in the larvae of *Drosophila*^84^ and the moth, *Manduca sexta* ^85,86^, but their physiology, downstream connectivity, and behavioral function had not been previously explored in adult flies.

We found that abdominal md neurons in the adult fly respond robustly to noxious heat (40°C) and a *TrpA1* agonist, as has been previously observed for the same cells in the larva. Our data suggest that detection of thermal nociceptive stimuli is conserved in these cells across metamorphosis **(Figure 3, Figure S3).** However, unlike in the larva^26,52,87^, we did not observe md neuron calcium signals in response to mechanical deflection of the abdomen. We also did not observe consistent changes in md axon activity when we stretched, squashed, or punctured the abdomen with a glass pipette (*data not shown*)—stimuli that have been shown to evoke calcium activity in larval class IV da neurons. Larval class IV da neurons are sensitive to shear forces^88^ and other mechanical stimuli that deform the larval body wall. Recent work has demonstrated that larval class IV da neurons can be activated by diffusible signals released from damaged tissue^89^, which may produce responses on longer timescales than we tested in our calcium imaging experiments. While we cannot rule out sensitivity to other mechanical stimuli, we did not observe responses to indentation or puncture of the abdomen in the adult fly. Their sensitivity to AITC suggests that adult md neurons retain *TrpA1* expression, and while *TrpA1* has been implicated in mechanosensation in other *Drosophila* sensory neurons^23^, our results suggest that md neurons in the adult fly primarily function as thermal nociceptors. More work is needed to understand the natural contexts within which abdominal md neurons are active and how they contribute to escape and sustained avoidance.

Shifts in sensory tuning from larva to adult may reflect adaptation to different ecological challenges. Adult flies do not burrow into the substrate and are capable of rapidly escaping into the air to evade predators. Ground-dwelling larvae, on the other hand, encounter distinct threats during burrowing and feeding, including parasitism^90,91^. The behavioral responses we observed in adult flies—jumping, running, and place avoidance—are more elaborate than the stereotyped rolling responses characteristic of larvae. This expanded repertoire may reflect the adult fly’s integration of nociceptive input with descending motor programs. We found that headless flies exhibited reflexive scratching and kicking, behaviors that may be suppressed by descending signals that drive escape.

In larvae, class IV da neurons connect to second-order interneurons that coordinate nociceptive responses. The most extensively studied are the Basin neurons, ascending interneurons that receive direct synaptic input from multiple md neuron types across body segments. Basin neurons are both necessary and sufficient for larval rolling escape responses^92–94^. Other pathways downstream of larval class IV da neurons include local circuits that coordinate segmental motor responses and descending neurons that modulate the magnitude of escape behaviors^30,95–97^. Unlike the organization we found in the adult **(Figure 5)**, larval class IV da neurons show less connectivity bias toward ascending pathways, with substantial connectivity to local motor circuits that drive the stereotyped rolling response. It is not clear whether Basin neurons and other second-order nociceptive interneurons survive metamorphosis^32,98^. Reorganization of second-order pathways may contribute to the expanded behavioral repertoire we observed in adults, as the larval circuits were optimized for the rolling escape response rather than the jumping, running, and avoidance behaviors seen in adult flies.

### Ascending Nociceptive Pathways

We found multiple ascending pathways positioned to transmit abdominal nociceptive signals to the brain **(Figure 6A–F).** Optogenetic activation of neurons labeled by SS51024, which connect to the giant fiber and other escape circuits, produced immediate reflexive escape without producing place avoidance (**Figure S6A,G**). In contrast, SS01159 neurons drove locomotion and sustained place avoidance; ascending cells within this line connect to the lateral horn^99–101^, dopaminergic PAM neurons, and MBONs, supporting their role in forming associative memories of dangerous locations (**Figure S6C,S6I**). IS35283 neurons produced an intermediate phenotype: smooth locomotor changes without strong jumping or avoidance (**Figure S6B,S6H**). While these ascending neuron classes recapitulate key features of md neuron activation, we note that our experiments relied solely on optogenetic activation. Future work could use optogenetic silencing to test the necessity of these pathways for nociceptive behaviors, or recordings of ascending neuron activity during natural stimulation.

The sustained behavioral effects we observed following md neuron activation, including place avoidance lasting at least 30 seconds, suggest that nociceptor activation may trigger longer-lasting changes in internal state beyond immediate motor responses. These state changes could involve neuromodulatory^102–105^ systems that alter the fly’s behavioral priorities, shifting from exploration to heightened vigilance. While our analyses focused on ascending excitatory pathways, the VNC circuits also contain inhibitory and neuromodulatory neurons that could contribute to long-lasting changes in internal state. The md sensory neurons may also release neuromodulators, as we observed high densities of dense-core vesicles in their synaptic terminals (**Figure S6E**). Neurotransmitter prediction algorithms also suggest that some of the ascending neurons labeled by IS35283 co-release serotonin along with acetylcholine^96,106^. These observations suggest that nociceptors and downstream neurons rely on neuromodulation to sustain altered behavioral states beyond the duration of the initial sensory stimulus.

Our connectomic analyses revealed that md neurons predominantly receive excitatory cholinergic input from other sensory neurons **(Figure S5A–C)**, including input from other sensory neurons (leg and abdominal bristles that sense innocuous touch, taste bristles, and other unknown sensory types**; Figure S5E)**. This lateral excitatory connectivity within the nociceptive system may amplify sensitivity^107–109^ when noxious stimuli activate multiple classes of exteroceptive somatosensory neurons. Direct input from tactile bristle neurons to md neurons suggests that touch could enhance nociceptive responses under certain conditions, as occurs during tactile allodynia in vertebrates (**Figure S5E**). Further work is needed to test the role of tactile input and excitatory feedback to md neurons in the adult fly, particularly in response to combined thermal and tactile stimuli or following tissue injury.

## Summary

Our findings reveal that adult *Drosophila* satisfy several of the criteria^10,110^ commonly used to define the experience of pain: dedicated nociceptors, ascending pathways connecting peripheral sensors to integrative brain centers, and a behavioral capacity for long-term avoidance of nociceptive stimuli. The ability to trace genetically-defined neural circuits at synaptic resolution makes the fly a powerful system for dissecting the neural circuits and computations that underlie nociceptive behaviors. The identification of ascending nociceptive pathways opens the door to understanding how these signals are used by brain circuits to guide navigation, learning, and action selection. Understanding how evolution has sculpted diverse nervous systems to balance protecting the body and behavioral flexibility may inform approaches to understanding and treating pain-related disorders.

## Acknowledgements and Support

We thank members of the Tuthill Lab for technical assistance and feedback on the manuscript. We thank the following people for providing feedback on the manuscript, reagents, equipment, and scientific guidance: Jay Parrish, Sama Ahmed, Wes Grueber, Ishmail Abdus-Saboor, Ellen Lesser, Gabrielle Sterne, Stefanie Hampel, Andrew Seeds, Kazuo Emoto, William Joiner, David Shepherd, and James Truman.

J.M.J. was supported by an HHMI Gilliam Fellowship and NIH T32 Predoctoral Training Program in the Neurosciences. Other support was provided by National Institutes of Health grants R01NS102333, R01NS128785, and U19NS104655, a Searle Scholar Award, a Klingenstein-Simons Fellowship, a Pew Biomedical Scholar Award, a McKnight Scholar Award, a Sloan Research Fellowship, the New York Stem Cell Foundation, and a UW Innovation Award to J.C.T., J.C.T is a New York Stem Cell Foundation – Robertson Investigator.

## Author Contributions

J.M.J. and J.C.T. conceived the study and wrote the manuscript. J.M.J. analyzed the connectivity, calcium imaging, and optogenetic datasets. A.S. conducted SPARC experiments and created the gorgeous confocal images of the fly abdomen and VNCs. A.M. collected and processed calcium imaging data from md axons. G.M.C. and S.W.B. collected optogenetic activation data in tethered flies. A.P.C. proofread the md axons and connected neurons in FANC and helped annotate neurons in the BANC.

## Lead contact

Further information and requests for resources and reagents should be directed to and will be fulfilled by the lead contact, John C. Tuthill.

## Data and Code Availability

Data is available on Dryad (10.5061/dryad.fbg79cp8k). Code for analyzing and visualizing md neuron connectivity in the EM dataset, calcium activity of md neurons, preference and velocity behavior during optogenetic experiments, and walking kinematics is located on GitHub (https://github.com/jesmjones/nociceptive_pathways_paper).

## Methods

### Key Resources Table

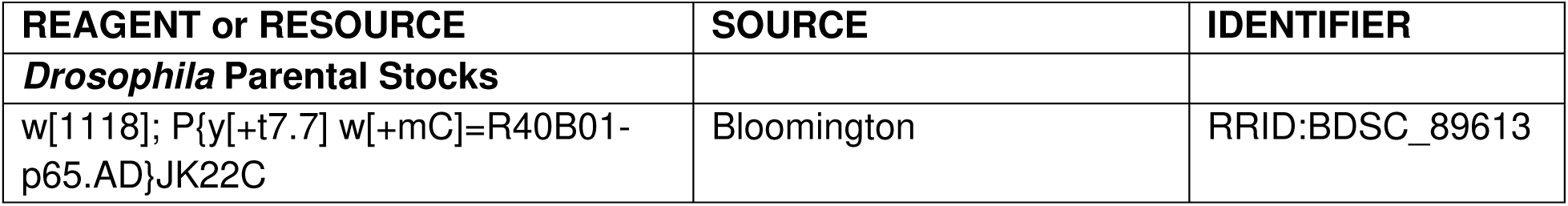
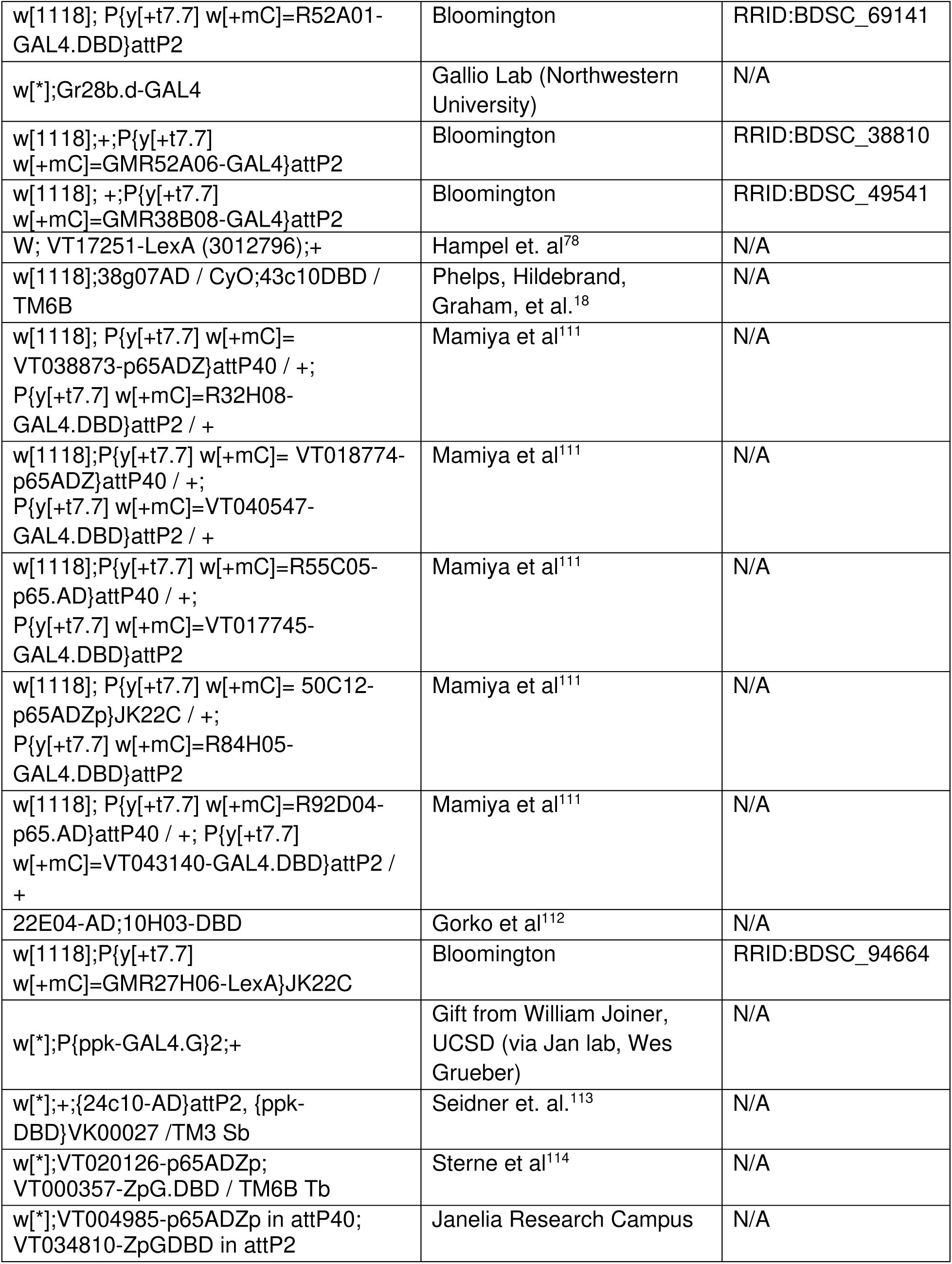
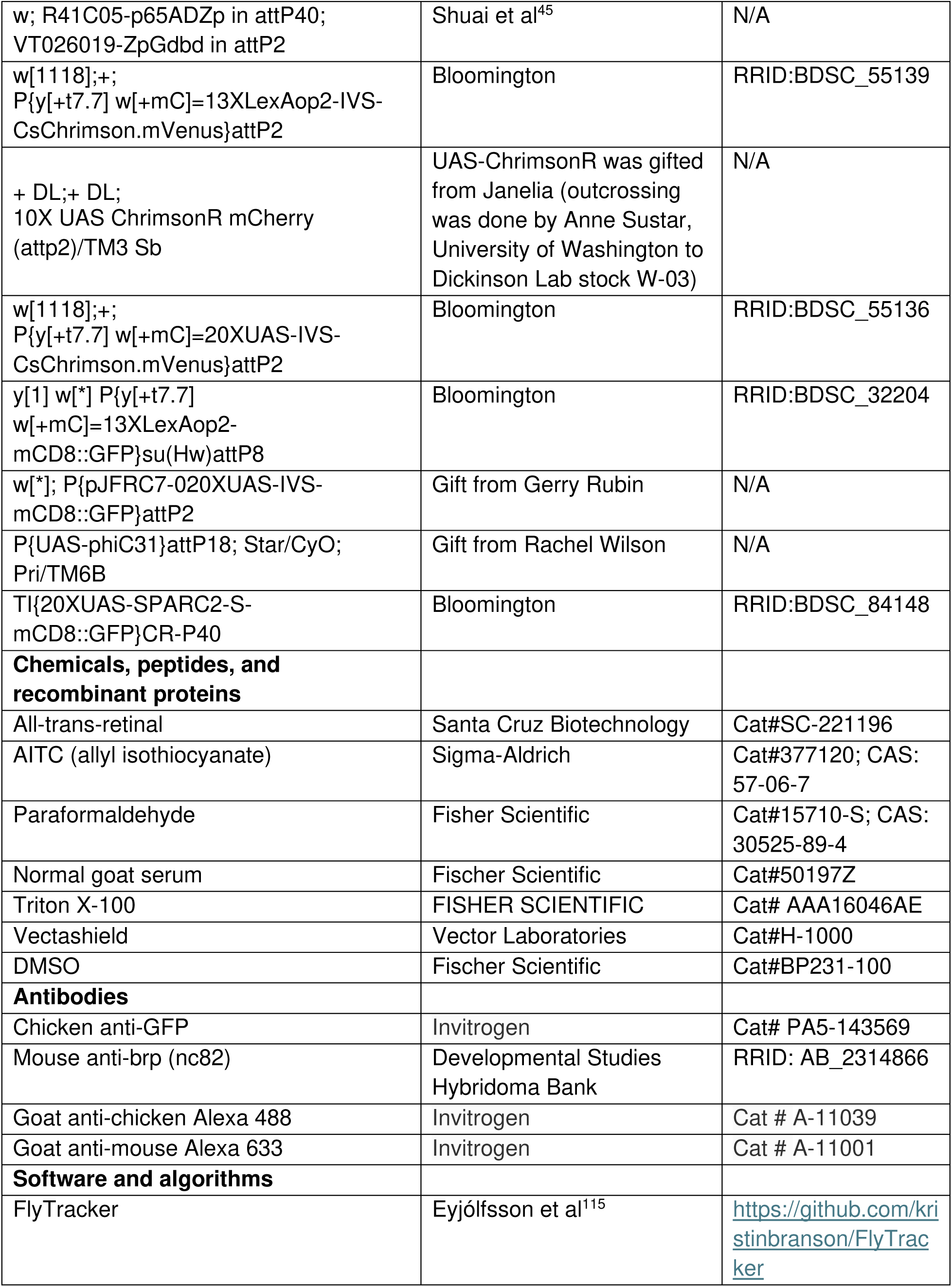
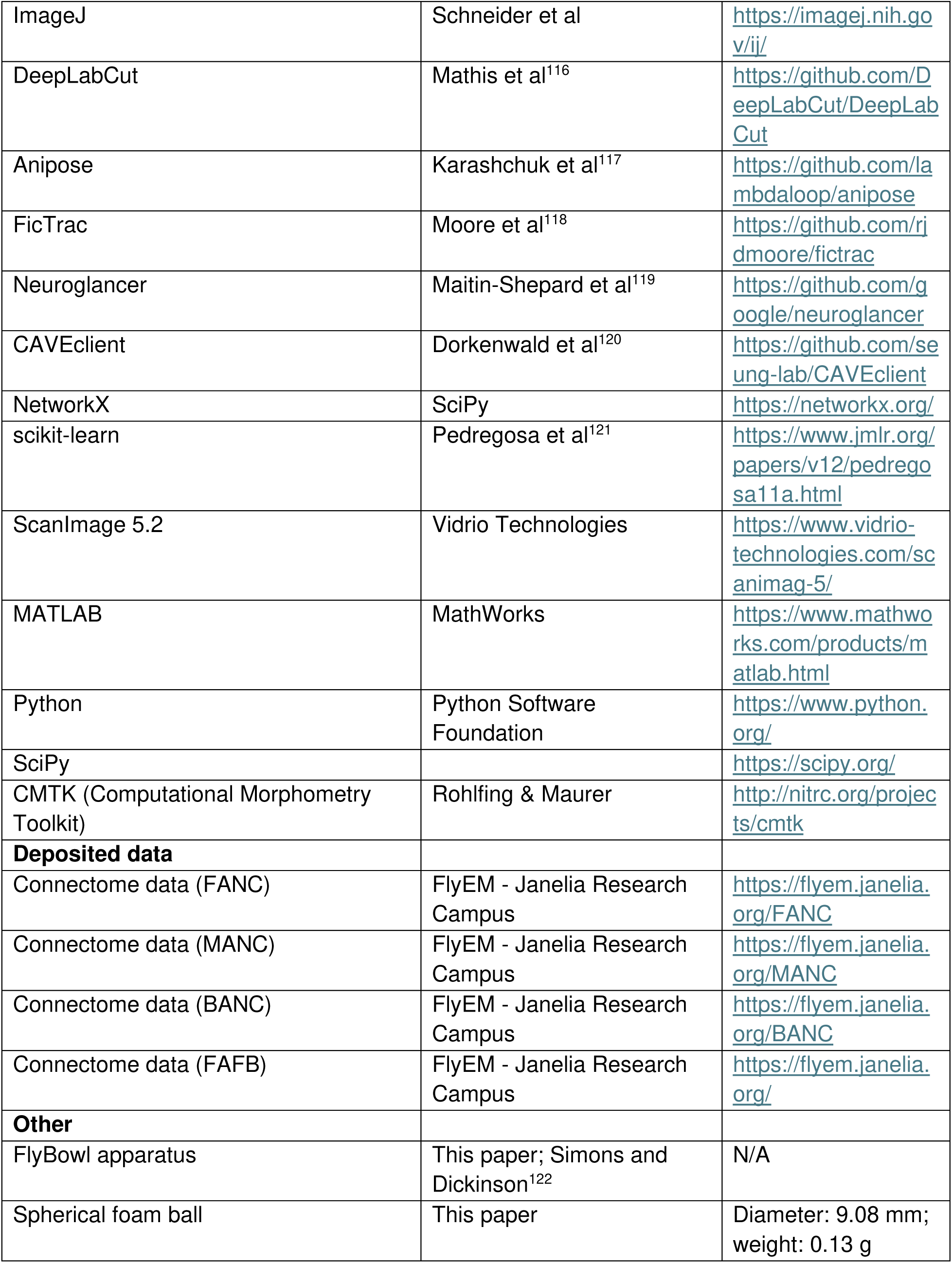

### Experimental Animals

All *Drosophila melanogaster* used in this study were raised on standard cornmeal molasses food and housed in an incubator kept at 25°C on a 14:10 light dark cycle.

### *Drosophila* Genotypes Figure Table

*Drosophila* Genotypes Figure Table:
| Figure Identifier | Full Genotype | Figure(s) |
| --- | --- | --- |
| 40B01-AD<br>(control) | w[1118]; P{y[+t7.7] w[+mC]=R40B01-p65.AD}JK22C > w[1118];+; P{y[+t7.7] w[+mC]=20XUAS-IV S-CsChrimson.mVenus}attP2 | 1C–E,<br>6A–C,E–<br>F |
| 52A01-DBD<br>(control) | w[1118]; P{y[+t7.7] w[+mC]=R52A01-GAL4.DBD}attP2 > + DL;+ DL; 10X UAS ChrimsonR mCherry (attp2)/TM3 Sb | 2B–C |
| hot cell | w[*];HC-GAL4;10X-UAS-ChrimsonR-mCherry (attp2) | 1C |
| b(A) | w[1118];+; P{y[+t7.7] w[+mC]=GMR52A06-GAL4}attP2/10X UAS ChrimsonR mCherry (attp2) | 1C |
| b(B) | w[1118]; +; P{y[+t7.7] w[+mC]=GMR38B08-GAL4}attP2/10X UAS ChrimsonR mCherry (attp2) | 1C |
| b(C) | W; VT017251-LexA (3012796) ; P{y[+t7.7] w[+mC]=13XLexAop2-IVS-CsChrimson.mVenus}attP2 | 1C |
| cs(A) | w[1118];38g07AD;43c10DBD/10X UAS ChrimsonR mCherry (attp2) | 1C |
| cho(B) (hook flexion) | W;VT038873-p65ADZ in attP40;32H08-DBD/10X UAS ChrimsonR mCherry (attp2) | 1C |
| cho(F) (hook extension) | W;VT018774-p65ADZ in attP40; VT040547-DBD/10X UAS ChrimsonR mCherry (attp2) | 1C |
| cho(A) (claw extension) | W;55C05 AD;VT017745 DBD/10X UAS ChrimsonR mCherry (attp2) | 1C |
| cho(D) (club) | w[1118]; P{y[+t7.7] w[+mC]= 50C12-p65ADZp}JK22C / +; P{y[+t7.7] w[+mC]=R84H05-GAL4.DBD}attP2/10X UAS ChrimsonR mCherry (attp2) | 1C |
| cho(C) (claw flexion) | W;92D04 AD;VT006391 DBD/10X UAS ChrimsonR mCherry (attp2) | 1C |
| cho(E) (neck chordotonal) | W ;22E04-AD;10H03-DBD/10X UAS ChrimsonR mCherry (attp2) | 1C |
| md(A) | w[1118];P{y[+t7.7] w[+mC]=GMR27H06-LexA}JK22C/P{y[+t7.7] w[+mC]=13XLexAop2-IV S-CsChrimson.mVenus}attP2 | 1C |
| md(C) | w[*];P{ppk-GAL4.G}2; P{y[+t7.7] w[+mC]=20XUAS-IV S-CsChrimson.mVenus}attP2 | 1C–E |
| md(B) | w[*];+;{24c10-AD}attP2; {ppk-DBD}VK00027/ P{y[+t7.7] w[+mC]=13XLexAop2-IV S-CsChrimson.mVenus}attP2 | 1C |
| md(B)>Chrimson | w[*];+;{24c10-AD}attP2; {ppk-DBD}VK00027 ;10X UAS ChrimsonR mCherry (attp2) | 2B–G |
| md(B)>GFP | w[*];+;{24c10-AD}attP2/ P{pJFRC7-02 20XUAS-IVS-mCD8::GFP}attP2; {ppk-DBD}VK00027 | S1H |
| md(C)>GFP | w[*];P{ppk-GAL4.G}2/P{pJFRC7-02 20XUAS-IVS-mCD8::GFP}attP2 | 3A, S1I |
| md(C)+ (control) | w[*];P{ppk-GAL4.G}2 | S1A–B |
| md(C)> CsChrimson | w[*];P{ppk-GAL4.G}2; P{y[+t7.7] w[+mC]=20XUAS-IV S-CsChrimson.mVenus}attP2 | S1A–E |
| md(A)>GFP | y[1] w[*] P{y[+t7.7] w[+mC]=13XLexAop2-mCD8::GFP}su(Hw)attP8;P{y[+t7.7] w[+mC]=GMR27H06-LexA}JK22C | S1F |
| IS35283>Chrimson | w[*];VT020126-p65ADZp; VT000357-ZpG.DBD / 10X UAS ChrimsonR mCherry (attP2) | 6A–F |
| SS51024>Chrimson | w[*];VT004985-p65ADZp in attP40; VT034810-ZpGDBD in attP2 / 10X UAS ChrimsonR mCherry (attP2) | 6A–F |
| SS01159>Chrimson | w; R41C05-p65ADZp in attP40; VT026019-ZpGdbd in attP2 / 10X UAS ChrimsonR mCherry (attP2) | 6A–F |
| SPARC | w[*];+;{24c10-AD}attP2/P{pJFRC7-02 20XUAS-IVS-mCD8::GFP}attP2 ;{ppk-DBD}VK00027 | S4C,E |
| SPARC | P{UAS-phiC31}attP18; Star/{24c10-AD}attP2; Pri/{ppk-DBD}VK00027 | S4A–B, S4D–G |
| SPARC | TI{20XUAS-SPARC2-S-mCD8::GFP}CR-P40; 24c10-AD}attP2; {ppk-DBD}VK00027 | S4A–B, S4D–G |
| md(C)-GAL4>UAS-GCaMP7f-tdTomato | w[*];P{ppk-GAL4.G}2/P{w[+mC] = UAS-tdTom.S}2;PBac{y[+t7.7] w[+mC]=20XUAS-IVS-jGCaMP7f}VK00005 | 3B–E |
| md(C)-GAL4>UAS-GCaMP7f-tdTomato | w[*];P{ppk-GAL4.G}2/P{w[+mC] = UAS-tdTom.S}2; PBac{y[+t7.7] w[+mC]=20XUAS-IVS-jGCaMP7f}VK00005 | S3 |

As in past studies^45,123,124^, we used individual AD or DBD split-GAL4 lines crossed to various UAS constructs as genetic controls. These split-half lines have the same insertion sites and genetic background as the full split-GAL4 driver lines but do not produce a functional GAL4 or effector expression without the complementary split-half. We prefer this approach, and have used it extensively in past work, because other commonly used “empty-GAL4” lines exhibit off-target expression in the VNC^125^. We also note that different Chrimson variants (ChrimsonR vs. CsChrimson) were used for different experiments due to genetic constraints (i.e., availability of LexA/GAL4 constructs, chromosomes, insertion sites, etc.). We did not observe any differences in behavior produced by different Chrimson variants.

### Statistical Analysis

We performed statistical comparisons between two groups using paired or unpaired two-tailed Student’s *t*-tests. We assumed data were normally distributed with similar variances across groups. We defined statistical significance as *p* ≤ 0.01, and denoted it by stars in the relevant figure panels. We conducted statistical analyses with Python (SciPy).

### Immunohistochemistry and Imaging of VNCs and abdomens

For confocal imaging of mcd8::GFP-labeled neurons in the VNCs, we dissected the VNC from 2-day old female adults in PBS. We fixed the VNC in a 4% paraformaldehyde PBS solution for 20 min and then rinsed the VNC in PBS three times. We put the VNC in blocking solution (5% normal goat serum in PBST) for 20 min, then incubated it with a solution of primary antibodies (chicken anti-GFP antibody, 1:50; anti-Brp mouse for neuropil staining, 1:50) in blocking solution for 24 hours at room temperature. At the end of the first incubation, we washed the VNC with PBS with 0.2% Triton-X (PBST) three times over two hours, then incubated the VNC in a solution of secondary antibody (anti-chicken-Alexa 488, 1:250; anti-mouse-Alexa 633, 1:250) dissolved in blocking solution for 24 hours at room temperature. Finally, we washed the VNC in PBST three times, once in PBS, and then mounted it on a slide with Vectashield (Vector Laboratories). We acquired z-stacks of each VNC on a confocal microscope (Olympus FV1000). We aligned the morphology of the VNC to a female VNC template in ImageJ with the Computational Morphometry Toolkit plugin (CMTK32;http://nitrc.org/projects/cmtk).

### SPARC labeling

To obtain the data in Figure S4, we combined a split-GAL4 driver (24c10-AD; ppk-DBD) with a PhiC31-based SPARC^59^ cassette (UAS-SPARC-STOP-GFP) and a PhiC31 recombinase source (UAS-PhiC31) for sparse labeling of neurons. PhiC31 recombination irreversibly removes/inverts the STOP cassette in a stochastic subset of GAL4+ cells, which permits UAS-driven GFP expression only in recombined cells. For each experimental fly, we waited 3-7 days after eclosion, dissected the abdomen and VNC, and kept them as pairs. We fixed and stained the VNCs as described above. Abdomens were fixed and stained and mounted the same way, but before staining, we cut the abdomens along the dorsal midline with dissecting scissors to open the abdomen into a single sheet, removing gut and ovaries, and being careful not to disturb the epithelial area. We acquired z-stacks of abdomens on a Leica DMI6000 widefield microscope (low magnification) and Olympus FV1000 (high magnification).

We scored each fly for (i) GFP expression in *ppk*+ neuron soma and dendritic morphology on the abdomen, and (ii) labeling density per hemisegment. For single-neuron analyses, we included animals with ≤2 GFP+ *ppk* neurons per hemisegment (preferably single-neuron). We excluded animals with dense labeling (>3 cells/hemisegment) from single-cell morphological and functional datasets.

### Optogenetics experiments in freely walking flies

For optogenetics experiments in freely walking flies (Figure 1), we housed adult flies on cornmeal-molasses food with dissolved all-trans-retinal (35 mM in 95% EtOH, Santa Cruz Biotechnology) and in the dark for at least 24 hr before experiments. We tested groups of 2-5 day old flies, separated by sex, in a 10 cm circular arena fitted with a glass top^122^. The arena was illuminated by infrared LEDs. For optogenetic activation, a red LED (625 nm-peak wavelength; ThorLabs) illuminated the arena from the top. Whole arena illumination experiments **(Figure 6C–E, Figure S1A–B)** had an LED intensity of 16.9 mW/mm^2^ in the center of the arena. Due to the placement of the LED, the quadrant arena experiments **(Figure 1**, **Figure 6B, 6F)** had a left LED intensity of 6.5-6.7mW/mm^2^, and a right LED intensity of 7.4-9.3mW/mm^2^. Boundary points, or edges of quadrants, had illuminations of 1.0mW/mm^2^ (left LED zone) and 1.4mW/mm^2^ (right LED zone). Off arena zones with no LED had an illumination of <0.1mW/mm^2^ during optogenetic activation. Fly behavior was recorded with a top-down view camera at 33 Hz (Basler acA1300-200 um; Basler AG) mounted with a lens (Computar). Videos were tracked using FlyTracker^115^, an automated system for tracking group walking trajectories and custom python script for analysis.

To quantify spatial aversion **(Figure 1**, **Figure 6B, 6F)**, we tracked flies in a circular arena divided into four equal quadrants. We designated two opposing quadrants as the stimulus zones, illuminating them with a red LED, while the remaining two quadrants served as control zones, with the LED off. At each time point, we classified the fly’s location as either within a stimulus (LED-on) zone or a control (LED-off) zone. We applied a binary scoring system: flies in the stimulus zones received a score of +1, and flies in the control zones received a score of –1. We then calculated a Preference Index (PI) for each fly. The resulting Preference Index ranges from –1 (complete avoidance of the LED zones) to +1 (complete preference for the LED zones), with 0 indicating no preference. We computed place preference statistics on a per-fly basis, across multiple trials.

**In Figure 1B**, to visualize spatial occupancy, we binned fly positions into a 2D histogram (3000 bins) for the time periods denoted by the trial (30 second bins), smoothed them with a Gaussian filter (sigma=50 pixels), and displayed them as a density heatmap where color intensity represents the frequency of occupancy at each location.

In **Figure S1,** we used a vector-based approach to analyze position data and quantify the directional consistency of fly trajectories during optogenetic stimulation trials. We grouped fly trajectory data by individual flies within each experimental trial. For each fly, we analyzed two predefined spatial regions of the arena: the *center* (the center 6cm of arena before the beveled edges) and the *surround* (the 2cm of border around the center that is beveled). We included only trials in which optogenetic stimulation was applied (opto condition of either ’left’ or ’right’) in the analysis. For each trial and arena region, we examined fly movement in two temporal windows: a **pre-stimulation** period (28–30 seconds) and a **post-stimulation** period (30–32 seconds). Within each period, we further analyzed time-series data to compute both instantaneous movement vectors and an overall **consistency** score. To compute movement vectors, we applied a sliding window across the trajectory data to extract incremental directional vectors between successive time points. We recorded each vector’s angle and magnitude to characterize local movement behavior. To quantify **directional consistency**, we used a custom function. This metric captures the degree to which the fly moved in a consistent direction over the two-second window. High consistency values (close to 1.0) reflect straight walking, whereas lower values (typically < 0.5) indicate more variable or turning behavior.

### Optogenetics experiments in tethered flies

We housed adult flies for optogenetics experiments in tethered flies on cornmeal-molasses food with dissolved all-trans-retinal (35 mM in 95% EtOH, Santa Cruz Biotechnology) and in the dark for at least 24 hr before experiments. We de-winged 2-5 day old flies and fixed them to a rigid tether (0.1 mm thin tungsten rod) with UV glue (KOA 300). We placed these flies onto a spherical foam ball (weight: 0.13 g; diameter: 9.08mm). We focused a red laser (638 nm; 1.2 kHz pulse rate; 30% duty cycle, Laserland) on the third segment of the abdomen (diameter of ∼350 µm). We conducted optogenetic activation experiments on flies in which md(B) flies (24c10AD; ppkDBD split GAL4) expressed ChrimsonR, as well as control flies. Trials were 2 seconds in duration and consisted of 500 milliseconds prestimulus, 1 second with the laser on, and 500 milliseconds post stimulus. During each trial, each fly was recorded with 6 high-speed cameras (300 fps; Basler acA800-510 um; Basler AG) and the movement of the ball was recorded at 30 fps with a separate camera (FMVU-03MTM-CS) and processed using FicTrac^118^. The 3D positions of each leg joint were determined using DeepLabCut^116^ and Anipose^117^. Kinematic analyses were performed with custom Python scripts.

### Analysis of behavioral data

**In Figure 2 and Figure S2,** to classify discrete motor behaviors in tethered flies, we analyzed femur-tibia joint angles recorded from the fly’s six legs over time. We used a threshold-based peak detection algorithm to identify significant movements within each joint’s trajectory. Specifically, we define peaks as local maxima in the joint angle time series that exceed a minimum height of 50 degrees and do not surpass 190 degrees. We chose these thresholds empirically to capture robust leg extensions while excluding small fluctuations or hyperextensions. We detected peaks separately for each leg and within each behavioral window.

We divided the behavioral data into consecutive, non-overlapping time windows of 35 frames. We chose this time window empirically to capture only one peak at a time. Within each window, we quantified which legs exhibit peak movement during the entire two-second trial (including the one-second stimulus period). We used this to classify the fly’s behavior during that window.

We classified behavioral states according to the specific pattern of detected peaks across legs:

- **Grooming** when only the hind legs (either left and right L3/R3 or L2/R2 midlegs) show rhythmic movement, while the other legs remain still. This pattern is consistent with grooming behaviors directed toward the head or body.
- **Kicking** when only a single leg shows movement during the window, typically indicative of isolated defensive or reflexive motions.
- **Standing** when no leg shows peak movement during each sliding time window.
- If none of the above conditions are met, we labeled the behavior as ambiguous or unclassified.

We applied behavior classification separately to each trial within the dataset. For each trial, we processed time windows sequentially, and we assigned classified behaviors to each frame in the original dataset.

To classify behaviors in the free-walking arena (**Figure 1D–E**, **Figure 6C**), we binned fly velocity (100 milliseconds). We classified flies with walking velocities less than 1mm/s as standing. We classified walking velocities between 1 and 30mm/s, which is the average top speed of flies walking in our arena, as walking. We classified flies that had large changes in x-y position as jumping. We applied these velocity bins to tethered flies to classify jumping and walking (**Figure 2D–E).**

We calculated the latency to jump following stimulus onset **(Figure 2E)** from annotated behavioral data for all trials. We filtered the data to include only frames occurring at or after the stimulus onset. We then grouped the dataset by trial and individual fly, and for each group, we identified the first occurrence of the "jumping" or "walking" behavior after stimulus onset. We calculated the latency to behavior as the difference between the frame number of the first behavioral event and the stimulus onset time. We assigned no latency value to trials in which no behavior occurred after stimulus onset.

To quantify the probability of behaviors over time (**Figure 2D–E),** we analyzed annotated behavioral data across individual flies. For each fly, we computed the number of frames labeled as a behavior and divided it by the total number of observed frames for that fly and time point. First, we counted all frames annotated as a given behavior for each fly and frame number. We then calculated total frame counts, regardless of behavior, for the same identifiers. We merged these counts by fly and frame number, and we computed the probability of the given behavior as the ratio of the number of frames for a given behavior to total time. We assigned a probability of zero for that time point to flies with no behavior annotations at a given frame.

In **Figure 2C**, we visualized heading angles of tethered walking flies using polar histograms to assess the distribution of movement orientations. We expressed behavioral trajectory data in degrees, normalized them to the range 0–360°, and converted them to radians for polar plotting. We binned angle values into 36 equal-width sectors (10° each) spanning 0 to 2π radians. We computed the frequency of headings within each bin and normalized them to represent proportions rather than raw counts. We generated all plots using Matplotlib in polar projection mode.

### Md and Postsynaptic Neuron Reconstruction in the connectome datasets

For the analyses in Figure 4, we first reconstructed md neurons and postsynaptic neurons in the female adult nerve cord (FANC^15^). We then matched sensory and postsynaptic partners in the male adult nerve cord (MANC^19^). We proofread automatically segmented neurons using Neuroglancer^119^, an interactive software for visualizing, editing, and annotating 3D volumetric data. Proofreading entailed two types of edits: we split off neurites that did not belong to the cell of interest, and we merged segments of the neuron that the automated segmentation falsely missed. Light-level images of md genetic driver lines guided our identification. We reconstructed md axons from abdominal nerves 2, 3, 4, and the fused posterior trunk. In FANC, the dorsal side of the abdominal ganglion was cut during dissection, which prevented us from following axons completely out of the nerve cord.

We used the reconstructed FANC neurons as a reference to identify homologous neurons in the MANC dataset. MANC neurons matched to FANC neurons shared the same postsynaptic connectivity. We based nerve assignment on both single axon labeling of peripheral neurons and reconstructed md neurons matched in FANC. Janelia proofreaders reconstructed axons in MANC, and further proofreading was not possible. For both FANC and MANC, we excluded the most posterior *ppk*+ neurons associated with the genitals (4–6 neurons) for all analyses.

Past work found that applying a 3-4 synapse threshold mitigates the inclusion of false positive connections^62,67,69^. For the analyses in Figure 5, we similarly analyzed postsynaptic neurons based on a synapse threshold of ≥4 synapses. In FANC, we reconstructed the top 100 neurons, sorted by the highest connectivity. We identified all objects in the automated segmentation that received ≥4 synapses from an md neuron. Synapses were detected automatically as described by Azevedo et al., 2024^15^. We then proofread those objects until they were associated with either a cell body or an identified descending or sensory process. We categorized a small number of objects as fragment segments, and we could not connect them to a cell body or an identified descending or sensory process. We deemed a neuron “proofread” once its cell body was attached, we reconstructed its full backbone, and we confidently attached as many branches as possible. Neuron annotations were managed by CAVE, the Connectome Annotation Versioning Engine. We used custom Python scripts to interact with CAVE via CAVEclient^120^. User authentication required to interface with CAVE related datasets.

We classified each md axon as either anterior or posterior based on whether the axon morphology branched anteriorly or posteriorly upon entering the VNC and which nerve bundle the axon traveled in. SPARC labeling confirmed the peripheral location of central md axons **(Figure S4).** We based the dorsal-ventral (DV) axes on SPARC labeling and python’s scikit-learn cosine_similarity function.

To compare md axons based on their synaptic connectivity in the insets of **Figure 4D–E**, we constructed cosine similarity matrices from filtered synaptic data using the scikit-learn python package. This process involved several stages:

1. **Filtering Raw Synapse Data:** We began by filtering the raw synapse dataset to retain only meaningful connections (≥4 synapses). For each presynaptic neuron, we identified all postsynaptic partners and counted the number of synapses between each pair. We retained only connections with a synapse count equal to or exceeding our defined 4 synapse threshold. This ensured that weak or potentially spurious connections did not contribute to the similarity analysis.
2. **Generating a Directed Weighted Graph:** From the filtered synapse data, we generated a directed weighted graph using NetworkX^126^. Each node represented a neuron, and each edge represented a synaptic connection, with weights corresponding to the number of synapses.
3. **Extracting a Connectivity Matrix:** We extracted a connectivity matrix where rows represented source (presynaptic) neurons and columns represented target (postsynaptic) neurons. For asymmetric analyses, the matrix included only directed edges from each presynaptic neuron to each postsynaptic target.
4. **Clustering Cosine Similarity Scores:** We then hierarchically clustered cosine similarity scores using the agglomerative clustering methods from the scikit-learn Python package.
5. **Sorting the Similarity Matrix:** To sort the similarity matrix, we applied agglomerative hierarchical clustering. We used the resulting dendrogram to reorder rows and columns, which allowed visualization of structurally related neuron groups. When analyzing asymmetric matrices, we clustered rows and columns independently based on their respective similarity profiles (i.e., dorsal to ventral, anterior to posterior).

### Analysis of central circuits downstream of md neurons

For the analyses in **Figure S6**, we constructed the jumping and walking connectivity networks for corresponding postsynaptic partners using data from MANC and BANC datasets via flywire.ai. We focused our tertiary analysis on described neurons in literature and neurons annotated by subcluster class in BANC and MANC. Having two datasets—one from the VNC alone and one from the entire nerve cord and brain—provides a more holistic view of circuits because some neurons only synapse in the brain, while others have connectivity in both the brain and nerve cord. We surveyed the broad connectivity of the ascending neurons, but for clarity chose to highlight specific connections to previously studied pathways that mediate behavioral phenotypes we observed in this study.

### Reconstructed neurons table

| Neurons | Dataset | Link |
| --- | --- | --- |
| md neurons | FANC | link |
| md neurons | MANC | link |
| md neurons | BANC | link |
| SS01159 | BANC | link |
| IS35283 lineage neurons | FANC | link |
| SS51024 | FANC | link |
| IS35283 lineage neurons | MANC | link |
| SS51024 | MANC | link |
| IS35283 lineage neurons | BANC | link |
| SS51024 | BANC | link |
| IS35283 brain axons | FAFB | link |
| SS51024 brain axons | FAFB | link |
| all putative aversive sensory neurons | MANC | link |
| all putative aversive sensory neurons | BANC | link |
| putative labellum *ppk*+ neurons brain axons | FAFB | link |

To analyze the connectivity of ascending neurons in each driver line, we cross-referenced each line to NeuronBridge to identify corresponding neurons in connectomic datasets. We queried postsynaptic targets of these neurons using three datasets: MANC (ventral nerve cord), FAFB (brain only), and BANC (brain and nerve cord combined). For each driver line, we identified ascending neurons based on morphological matching and analyzed their synaptic outputs to understand potential behavioral circuits.

For SS51024, we matched excitatory ascending neurons across datasets and analyzed their connectivity to the giant fibers, premotor networks, and brain circuits. For IS35283, we identified excitatory ascending neuron pairs and examined their connections to walking control circuits and descending neurons. For SS01159, we identified 60 ascending neurons (57 cholinergic, 3 glutamatergic based on predicted neurotransmitter classification) and focused analysis on the 6 neurons with strongest md connectivity (>5 synapses per neuron).

We quantified synaptic weights as total synapses between pre- and postsynaptic partners. We quantified VNC synapses in MANC, while we quantified brain synapses in FAFB. Cross-dataset analysis allowed us to integrate nerve cord and brain connectivity. We used BANC reconstructions for visualization of ascending neurons.

### *in vivo* two-photon calcium imaging

We used a two-photon Movable Objective Microscopes (MOM; Sutter Instruments) with a 40x water-immersion objective (0.8 NA, 2.0 mm wd; Nikon Instruments) for calcium imaging.

We used a mode-locked Ti:sapphire laser (Chameleon Vision S; Coherent) to excite fluorophores at 920 nm. We maintained power at the back aperture of the objective below ∼25 mW with a Pockels cell. Emitted fluorescence was directed to two high-sensitivity GaAsP photomultiplier tubes (Hamamatsu Photonics) through a 705 nm edge dichroic beamsplitter followed by a 580 nm edge image splitting dichroic beamsplitter (Semrock). Fluorescence was band-passed filtered by either a 525/50 (green) or 641/75 (red) emission filter (Semrock). Image acquisition was controlled with ScanImage 5.2^127^ (Vidrio Technologies) in MATLAB (MathWorks). The microscope was equipped with a galvo-resonant scanner, and the objective was mounted onto a piezo actuator (Physik Instrumente; digital piezo controller E-709). We focused our recordings on the abdominal ganglion, where the md axons project, using a fly holder and dissection similar to past work^128^. All experiments were performed in the dark at room temperature with 24C10AD;ppkDBD-GAL4>UAS-GCaMP7s-tdTomato flies. We used a 25W, 120V consumer soldering iron probe (Model SP25NKUS; Weller) attached to a 2000VA Auto Variable Voltage Transformer for temperature control (VEVOR). We calibrated the voltage necessary to achieve 40°C using a thermocouple attached to the iron tip and to a PID controller (Jaybva; PID temperature controller meter indicator).

In the experiments in **Figure 3**, we applied a 40°C heat or room temperature 25°C probe to dorsal abdominal segments 2-5. In **Figure S3C–E,** we applied 50 µM AITC in DMSO to the dorsal surface of the abdomen using a paintbrush. We pseudo-colored images to enhance the signal-to-noise ratio, where yellow represented the highest intensity and dark blue the lowest. We processed calcium imaging data to extract fluorescence signals and generate normalized activity traces for each experiment. We imported raw fluorescence data from tdTomato and GCaMP channels from individual CSV files and concatenated them into a single dataset. We parsed metadata such as recording date, fly identity, genotype, and trial number from filenames. For each recording, we calculated a fluorescence baseline (F₀) from the initial portion of the imaging period and computed ΔF/F values separately for the tdTomato and GCaMP channels. To account for motion or expression variability, we calculated the GCaMP-to-tdTomato fluorescence ratio for each frame and used its baseline to derive a normalized ΔR/R_0_ signal. We annotated stimulation periods by aligning fluorescence traces to a metadata table specifying the onset and offset frames of each heat stimulus. We labeled each frame as either “stimulus on” or “stimulus off,” and marked recordings identified as mechanical controls accordingly.

---

**Extended Data Figure S1:**
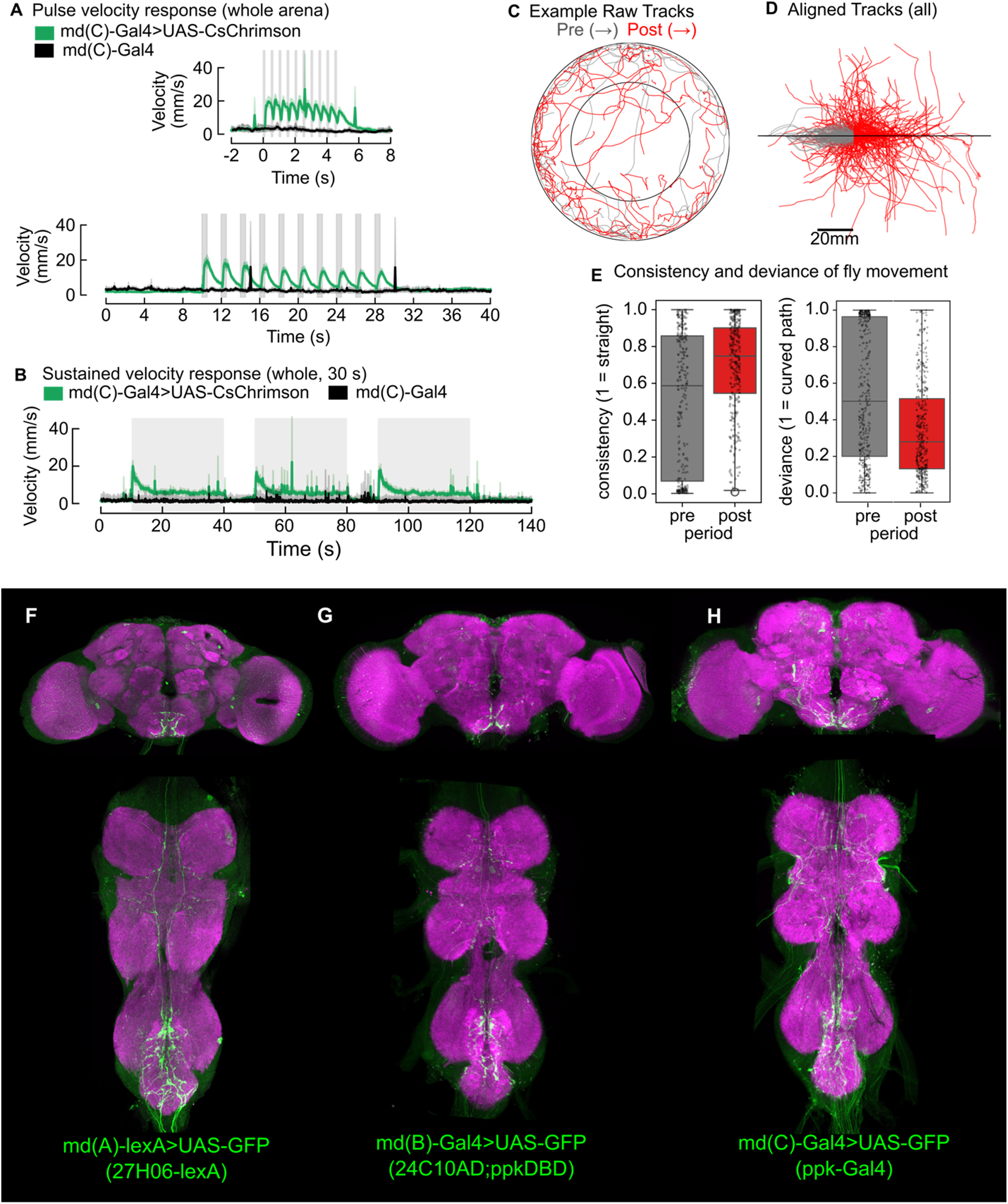
Pulse activation experiments of md neurons and driver lines used. (A-B) Repeated activation of md(C)-GAL4 expressing UAS-CsChrimson (A), 500ms (A), and 30 seconds (B). C-E) Directional quantification of flies in quadrant arena assay. (C) Example raw tracks of flies pre (grey) and post(red). (D) Aligned tracks of all flies. (E). Quantification of consistency and deviance of fly movement. See Methods for details. (F-H) Genetic driver lines used to label abdominal multidendritic neurons (Figure 1C). Each driver line expressed in abdominal md neurons and other (non-overlapping) cell-types.

**Extended Data Figure S2.**
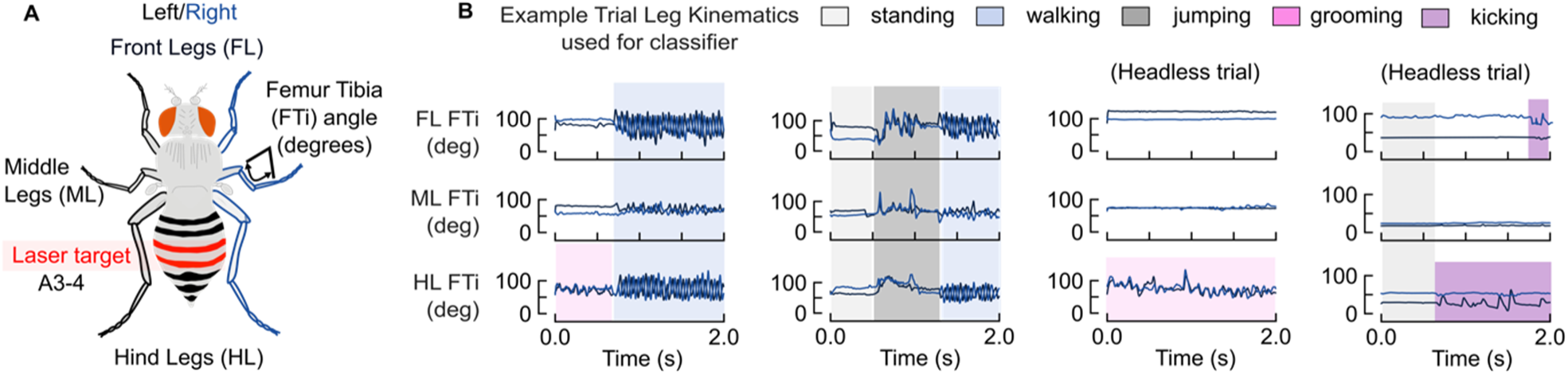
Kinematics used to perform behavioral classification of flies on the ball. (A) Schematic of fly with front legs (FL), middle legs (ML), and hind legs (HL), with example femur-tibia angle (FTi°) used to plot joint kinematics. (B) Plotting of leg FTi° over time with highlighted stretches to signify standing (white) walking (blue) jumping (dark grey) abdomen grooming (pink) and single leg kicks (purple).

**Extended Data Figure S3.**
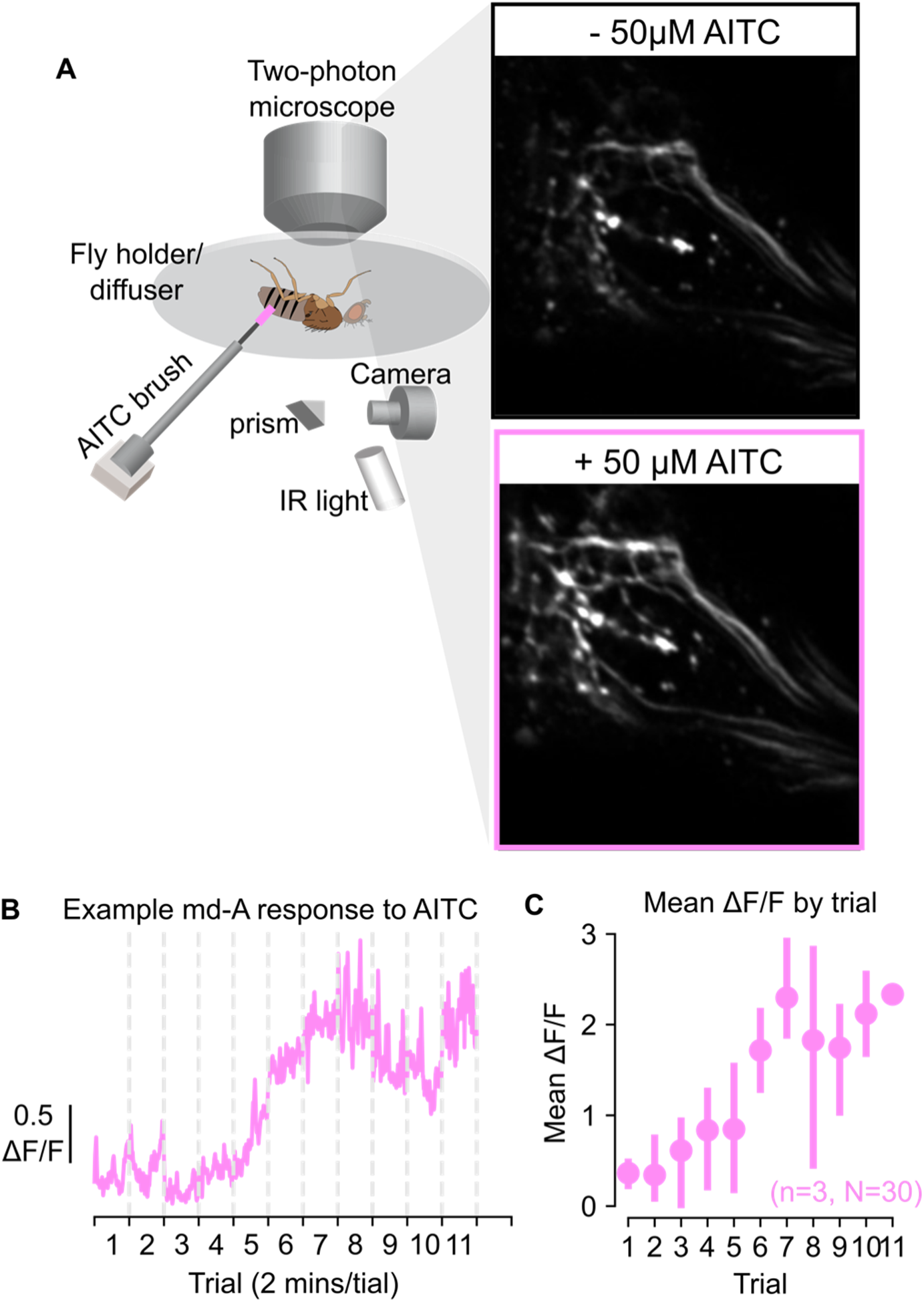
Application of AITC increases calcium activity in md axon terminals. (A) We administered 50 micromolar AITC to the abdominal wall using a paintbrush and quantified the change in fluorescence over multiple imaging trials. (B) All trials were normalized to Trial 1. (C) Mean fluorescence of each trial. N=30 trials total across n=3 flies

**Extended Data Figure S4.**
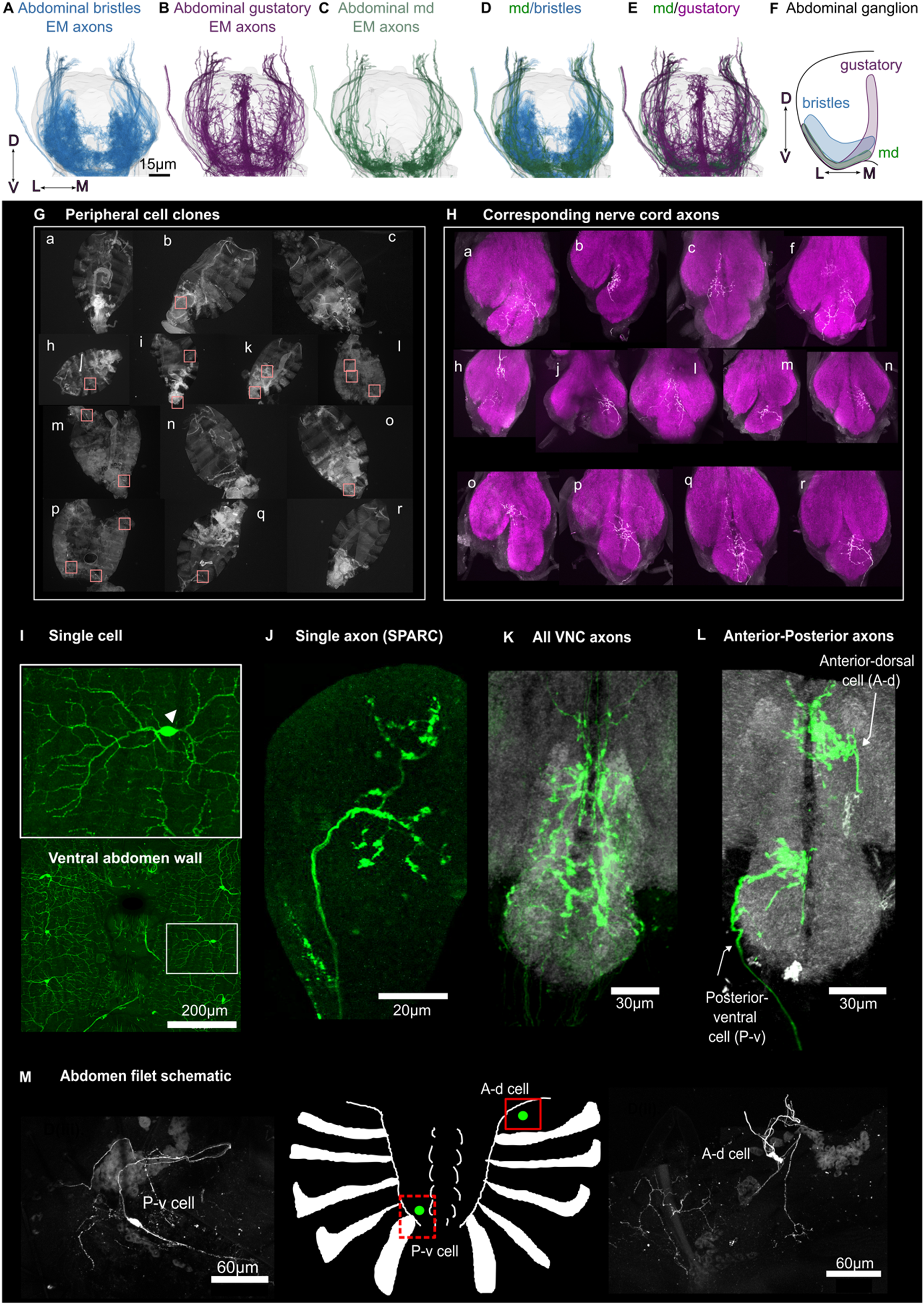
Neuropil domains and SPARC labeling of 24C10AD;ppkDBD-GAL4. (A-E) MANC Dorsal-ventral EM reconstruction views of abdominal bristle axons (blue, A), abdominal putative gustatory neurons (purple, B), abdominal md axons (green, C), md-bristle axons (green-blue, D), and md-gustatory axons (green-purple, E). Scale bar is 15 microns. View is of abdominal ganglion. (F) Neuropil domains of each of the three sensory axon subtypes in the abdominal ganglion. Gustatory and md axons are more ventral than bristle axons, and bristle axons span more of the ventral domain of the abdominal ganglion. (G-H) Peripheral cell clones and their corresponding VNC axons. Lowercase letters are matched between abdominal and vnc labeled axons. Pink boxes denote locations of cell bodies. (I) Morphology of a peripheral md cell labeled by GFP. Bounding box encompasses one cell and its receptive field. Triangle denotes location of cell body. J) Single axon of a SPARC clone in the VNC labeled by GFP. (K) All VNC axons labeled by 24C10AD;ppkDBD-GAL4>UAS-GFP. (L) Anterior-dorsal and posterior-ventral single cell clones from SPARC protocol. (M). Abdomen filet schematic, origin location, and confocal images of anterior-dorsal (A-d) and posterior-ventral (P-v) cells. Genotype used throughout SPARC labeling was md(B)-GAL4 (24C10AD ; ppkDBD). SPARC procedure outlined in Methods

**Extended Data Figure S5.**
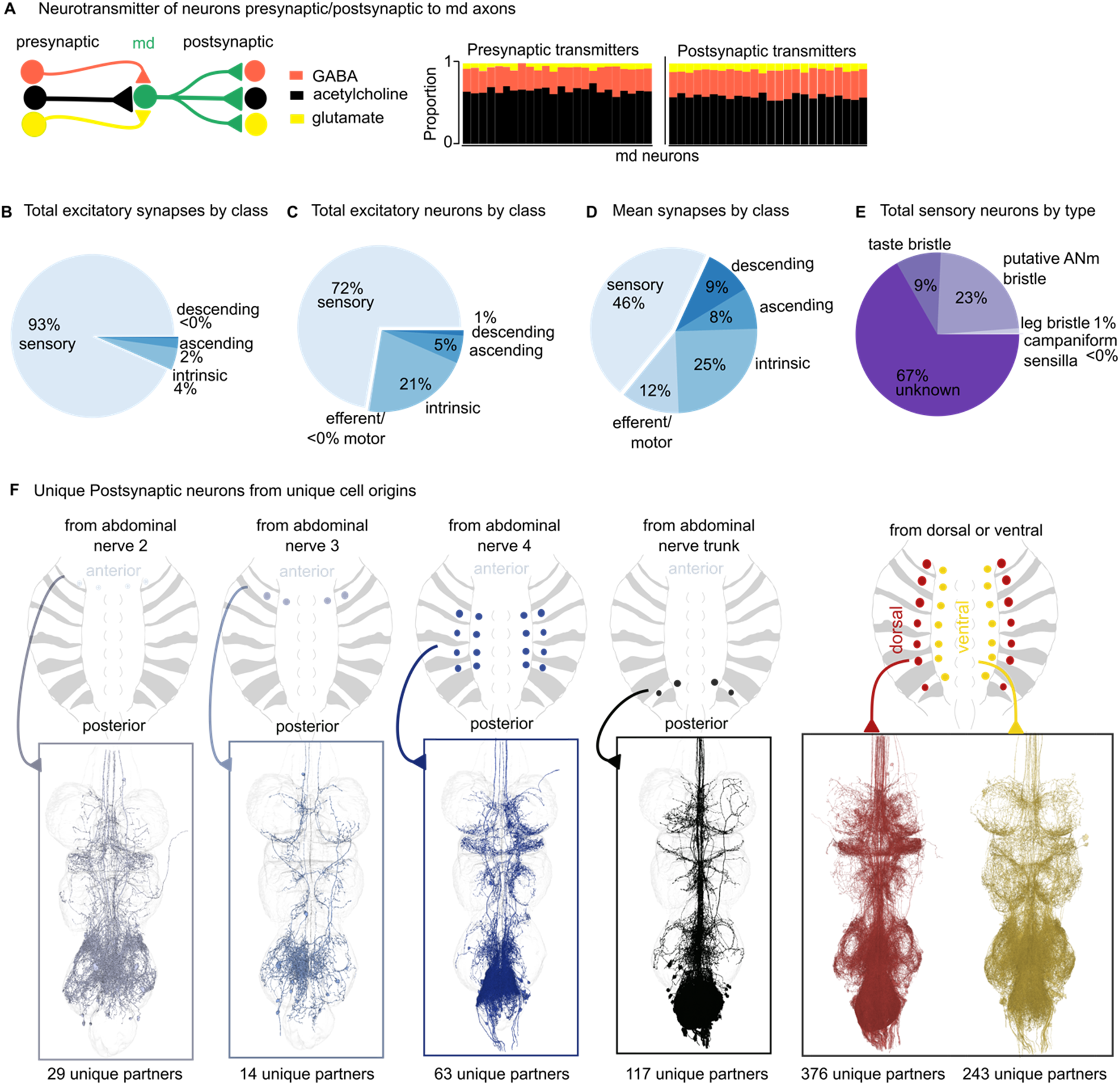
Connectivity and neurotransmitter analysis of abdominal md neurons. (A) Neurotransmitter identity of neurons presynaptic and postsynaptic to md axons. Md neurons predominantly receive excitatory cholinergic input and synapse onto cholinergic postsynaptic neurons. (B-D) Connectivity analysis of md neuron outputs by postsynaptic partner class. (B) Total excitatory synapses, (C) total excitatory neurons, and (D) mean synapses to md neurons. (E) Pie charts show sensory neurons by proportion presynaptic to md neurons (campaniform sensilla, leg bristles, putative ANm bristles, and taste bristles). (F) Spatial organization of md neuron connectivity by nerve of origin. Cross-sections of abdominal ganglion show postsynaptic partner distributions for md neurons entering via different nerves (nerve 2, 3, 4, and nerve trunk), with corresponding reconstructed axon morphologies below. Numbers indicate unique postsynaptic partners per nerve group. Md neurons originating from dorsal abdomen (red) and ventral abdomen (yellow) show distinct spatial distributions of their postsynaptic targets within the abdominal ganglion, preserving somatotopic organization. Cross-section shows target neuron locations with corresponding reconstructed md axon morphologies. Data from MANC connectome.

**Extended Data Figure S6.**
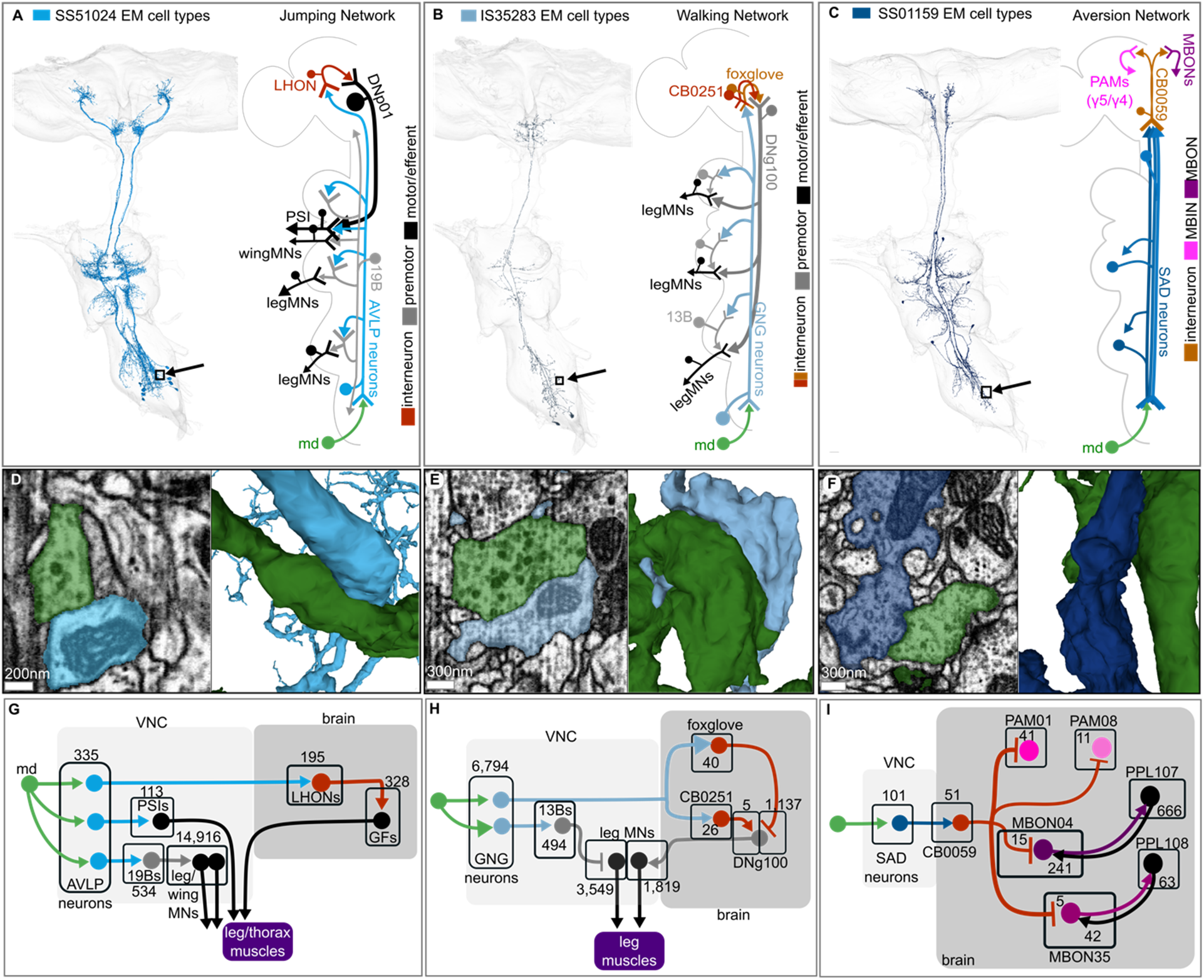
Connectivity of ascending neurons to known circuits in the brain. (A-C) Circuit diagrams showing connectivity between md-related ascending neurons and established central brain neurons and networks. (A) SS51024 jumping network. md neurons connect to ascending neurons that integrate with giant fiber (GF) escape circuits through peripherally synapsing interneurons (PSIs) and lateral horn output neurons (LHONs). Pathway includes connections to 19B premotor neurons targeting leg and wing motor neurons (MNs), and descending neuron DNp01 controlling takeoff behavior. (B) IS35283 walking network. md neurons connect to ascending neurons projecting to gnathal ganglion (GNG) that integrate with walking control circuits. Key connections include CB0251 and foxglove interneurons that synapse onto descending neuron DNg100, and 13B premotor neurons targeting leg motor neurons. (C) SS01159 aversion network. md neurons connect to ascending neurons projecting to multiple brain regions including saddle (SAD). Circuit includes connections through CB0059 interneurons to PAM dopaminergic clusters (γ4, γ5) that drive mushroom body input neurons (MBINs) and output neurons (MBONs) involved in aversive learning. We hypothesize that these are the neurons involved in the avoidance we see, but further work is needed to validate these neurons. (D-F) Electron microscopy images showing synaptic connections between circuit components at coordinates indicated by the box and arrows on EM reconstructions in A-C. Scale bars: 200-300 nm as indicated. (G-I) Subcircuits downstream of ascending neurons. Numbers in black boxes indicate synapse counts between connected neuron types. Connectomic analysis performed across MANC and FAFB datasets. Gray boxes indicate VNC vs. brain. See Methods for circuit reconstruction details.

